# Chronic *in vivo* optogenetic stimulation modulates neuronal excitability, spine morphology and Hebbian plasticity in the mouse hippocampus

**DOI:** 10.1101/320507

**Authors:** Thiago C. Moulin, Lyvia L. Petiz, Danielle Rayêe, Jessica Winne, Roberto G. Maia, Rafael V. Lima da Cruz, Olavo B. Amaral, Richardson N. Leão

## Abstract

Prolonged increases in excitation can trigger cell-wide homeostatic responses in neurons, altering membrane channels, promoting morphological changes and ultimately reducing synaptic weights. However, how synaptic downscaling interacts with classical forms of Hebbian plasticity is still unclear. In this study, we investigated whether chronic optogenetic stimulation of hippocampus CA1 pyramidal neurons in freely-moving mice could (a) cause morphological changes reminiscent of homeostatic scaling, (b) modulate synaptic currents that might compensate for chronic excitation, and (c) lead to alterations in Hebbian plasticity. After 24 h of stimulation with 15-ms blue light pulses every 90 s, dendritic spine density and area were reduced in the CA1 region of mice expressing channelrhodopsin-2 (ChR2) when compared to controls. This protocol also reduced the amplitude of mEPSCs for both the AMPA and NMDA components in *ex vivo* slices obtained from ChR2-expressing mice immediately after the end of stimulation. Lastly, chronic stimulation impaired the induction of LTP and facilitated that of LTD in these slices. Our results indicate that neuronal responses to prolonged network excitation can modulate subsequent Hebbian plasticity in the hippocampus.

## Introduction

Hebbian forms of synaptic plasticity, such as long-term potentiation (LTP) and long-term depression (LTD), are believed to be fundamental for learning and memory. However, by inducing long-lasting changes in synaptic strength, these mechanisms can drive circuit saturation and destabilization if left unchecked (Abbott & Nelson, 2000). As a counterpart to input-specific Hebbian plasticity, thus, neurons can sense changes in their excitability and initiate global homeostatic regulations in synapses to restrain their activity to its usual physiological range, in a process known as synaptic scaling (Turrigiano, 2012). These homeostatic processes act to ensure network stability, providing a feedback that prevents unrestrained potentiation or depotentiation stemming from LTP and LTD (Vitureira & Goda, 2013).

This interplay is classically mediated by scaling-induced changes in receptor density that compensate those brought upon by LTP and LTD; in this sense, however, homeostatic scaling could also affect subsequent induction of Hebbian plasticity itself. It has been proposed that synaptic scaling could modulate the threshold for LTP or LTD induction as a response to the changes in post-synaptic activity (Rabinowitch & Segev, 2008; Keck et al., 2017). This idea is supported by the fact that synaptic scaling and Hebbian plasticity, although operating under distinct plasticity rules, share several molecular pathways that converge to regulate synaptic morphology and membrane channel composition (Fernandes & Carvalho, 2016).

Moreover, there is experimental evidence that homeostatic plasticity induced by chronic neuronal inhibition can enhance subsequent LTP, as blockade of neuronal activity by tetrodotoxin (TTX) for 60 h in cultured mouse hippocampal slices was shown to increase the expression of AMPA and NMDA receptors at existing synapses. This prolonged inhibition was also able to create new synapses containing only NMDA receptors, which were recruited when LTP was induced at CA3-CA1 synapses, thus enhancing potentiation (Arendt et al., 2013). Another study demonstrated that acute deprivation of activity in hippocampal slices for only 3 hours by TTX and APV could also drive upscaling of spontaneous synaptic transmission. This brief period of homeostatic regulation was enough to facilitate LTP at CA3-CA1 synapses as well (Félix-Oliveira et al., 2014).

The effects of prolonged decreases in neuronal input can also be observed *in vivo*. Suppression of neuronal activity in the mouse primary visual cortex, caused by 2 days of visual deprivation, was able to promote LTP and prevent LTD in layer 2/3 pyramidal neurons (Guo et al., 2012), suggesting that this protocol, known to induce homeostatic upscaling (Goel et al., 2011), was sufficient to slide the threshold between synaptic potentiation and depression. A more recent study, however, has shown that this metaplastic effect depends on NMDA receptor activity rather than on neuronal firing alone (Bridi *et al.*, 2018), suggesting that it could be brought about by Hebbian plasticity rather than synaptic scaling.

Despite the evidence for enhancement of LTP by activity suppression, there is little data on the effects of chronic neuronal excitation (and putative synaptic downscaling) on Hebbian plasticity; nevertheless, there is reason to believe that these processes interact as well. As opposite processes, up- and downscaling differ in their need for signaling and trafficking elements, but both of them modulate mechanisms that underlie synaptic potentiation and depression, such as synapse morphology, glutamate transport and vesicle function (Turrigiano, 2012; Schanzenbächer et al., 2018). Furthermore, a recent study showed that 10 min of high-frequency optogenetic stimulation could induce homeostatic plasticity in organotypic hippocampal cultures, decreasing the number of excitatory spine synapses and increasing the number of inhibitory ones. This downscaling protocol impaired memory recall and facilitated memory extinction *in vivo* when selectively applied to granule cells encoding a contextual fear memory (Mendez et al., 2018), although its effects on Hebbian plasticity were not explored.

To unveil the interaction between homeostatic response to overexcitation and Hebbian plasticity, as well as to establish a way to study the effects of synaptic downscaling in *vivo*, we develop a model using chronic optogenetic stimulation of pyramidal cells in the hippocampal CA1 region of freely-moving C57BL/6 mice. We then evaluate the effect of this intervention on synaptic morphology, neuronal excitability and synaptic plasticity in *ex vivo* brain slices.

## Results and Discussion

We induced the expression of channelrhodopsin 2 (ChR2) in CA1 pyramidal neurons of the hippocampus by injecting a viral vector (AAV5-CaMKIIa-ChR2-eYFP). To verify opsin expression and functionality, we obtained *ex vivo* slices (Figure 1A-B) after 3 to 5 weeks and activated ChR2-expressing neurons by a blue light-emitting laser. Voltage-clamp recording was performed to verify the activation of these channels (Figure 1C). We also tested if a 15-ms pulse would enhance overall excitation in the CA1 circuitry. We chose this emission time because it would be theoretically sufficient for ChR2 to reach its maximum response (Lin et al., 2009) while minimizing neuronal damage caused by prolonged photostimulation. In extracellular recordings of local field potentials (LFPs) in the stratum pyramidale, the mean number of spikes detected increased around twofold in the 1-second period following stimulation (Figure 1D; paired t test, p=0.013, n=7 slices).

**Figure 1:**
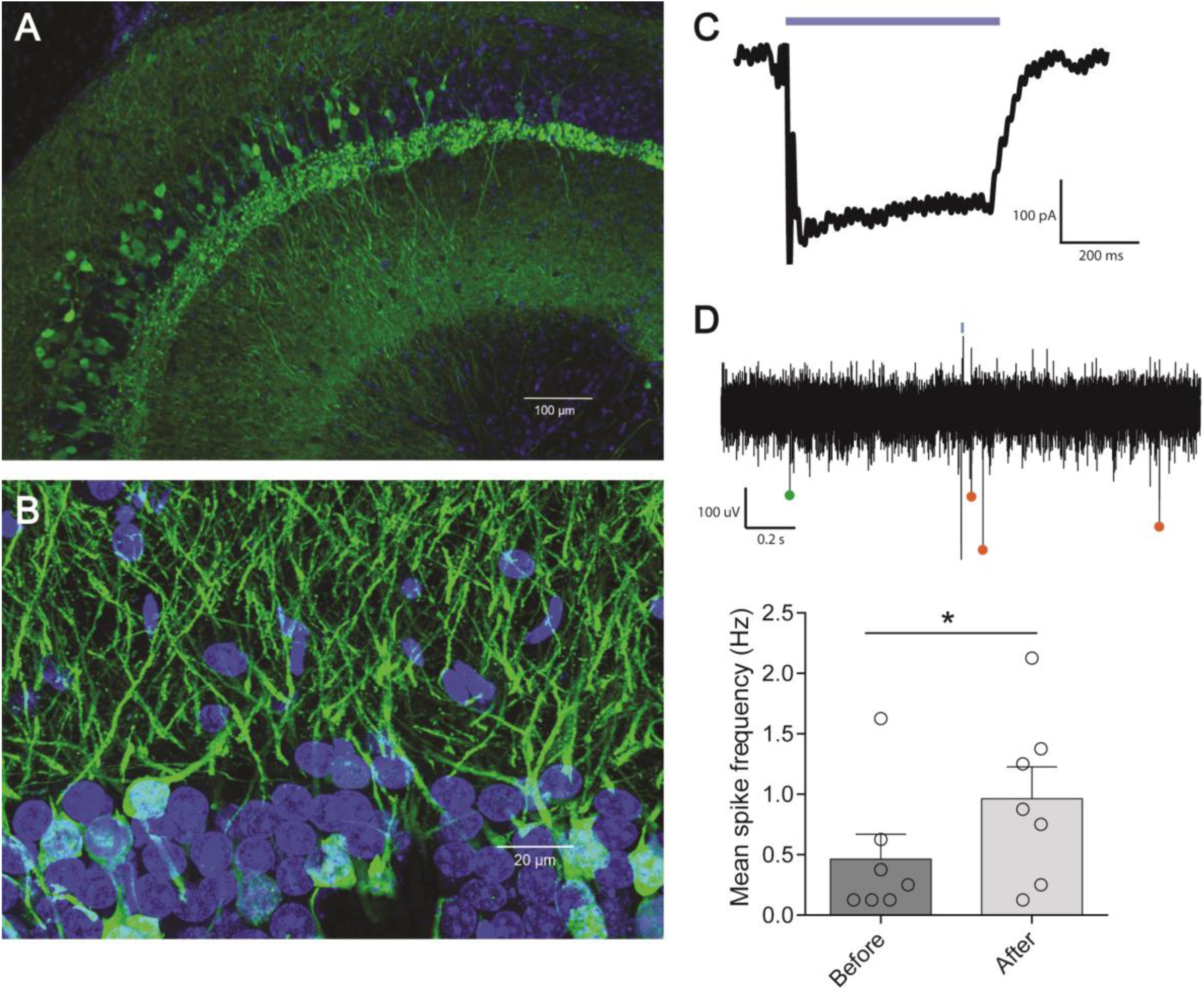
Expression of ChR2 in the hippocampus. (A) Overview of ChR2-eYFP expression in hippocampal slices. ChR2 is associated with eYFP (green) and expressed under the CaMKII-α promoter, while nuclei are stained in blue with DAPI. (B) Expression in pyramidal cells of the CA1 region. (C) Qualitative example of a voltage clamp recording of a neuron expressing ChR2 in response to 473-nm light. (D) Field activity recorded during 1-s intervals immediately before and after an acute 15-ms optogenetic pulse (shown as a blue line). Dots show examples of spikes detected before (green) and after (orange) stimulation in a recording. Bars show mean number of spikes (± s.e.m) before and after stimulation across 5 trials per slice, with circles showing the results for individual slices (n=7 slices; paired t test, p=0.013)

To test the effects of chronic *in vivo* optogenetic stimulation (Figure 2), we applied 15 ms of blue light every 90 s during 24 h through a fiber-optic cannula implanted in the CA1 region of freely-moving mice expressing either ChR2 or an eYFP-only viral vector (AAV5-CaMKIIa-eYFP). Immediately after chronic stimulation, mice were euthanized and hippocampal slices were used for either dendritic spine morphology assessment, mEPSC recordings or *ex vivo* induction of Hebbian plasticity.

**Figure 2:**
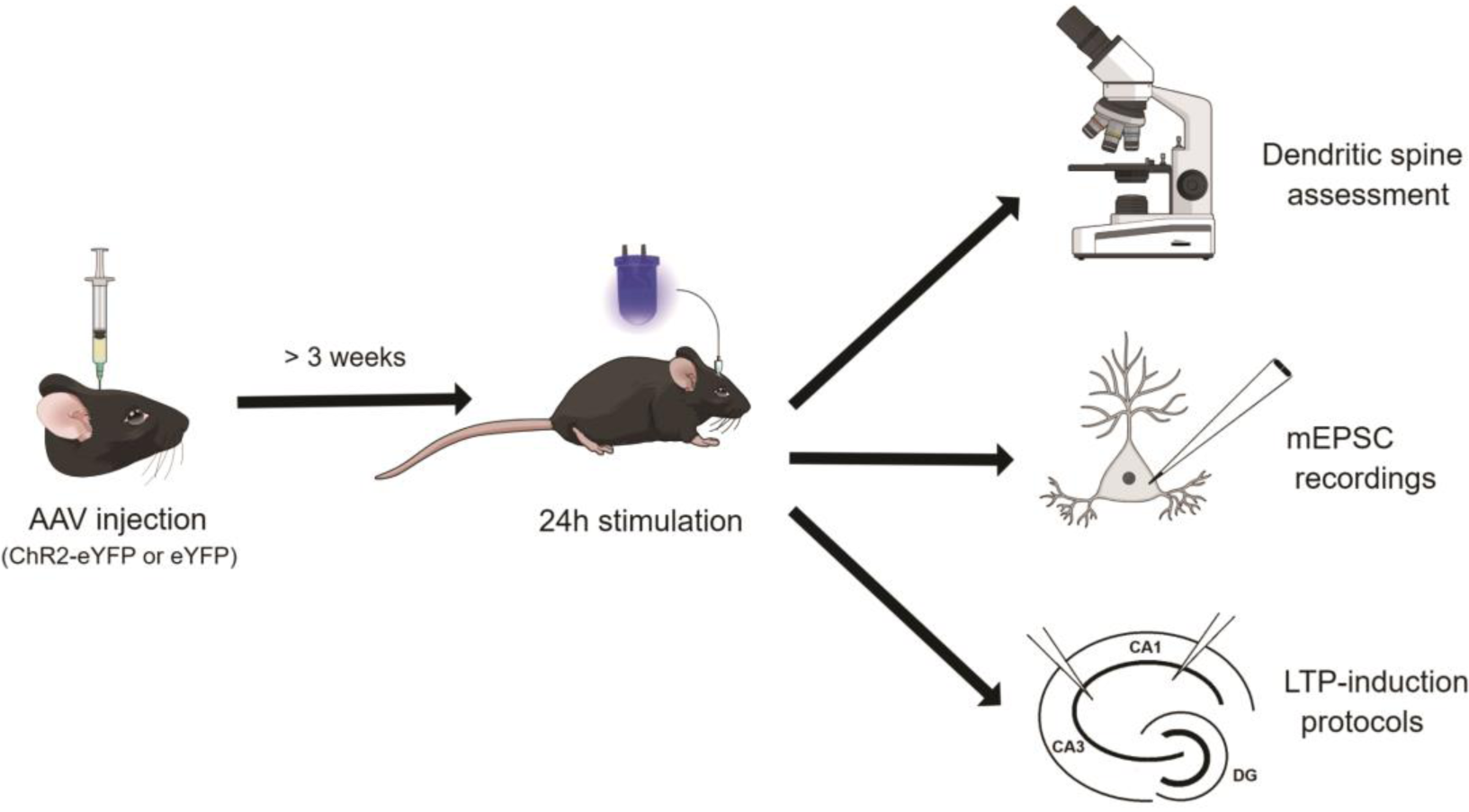
Experimental protocol. Three weeks after injecting a viral solution in the ventral hippocampus to induce the expression of ChR2-eYFP or eYFP alone, we applied 15-ms light stimuli through fiber-optic cannulae every 90 seconds during a 24-hour period. At the end of this period, hippocampal slices were either (a) fixed and processed in a confocal microscope after anti-GFP/YFP immunohistochemistry, (b) prepared for whole-cell patch-clamp to record miniature excitatory postsynaptic currents (mEPSCs), or (c) used for LTP induction protocols, with recording of field evoked post-synaptic potentials (fEPSPs) in the stratum radiatum of CA1 in response to Schaffer collateral stimulation.

To visualize dendritic spines, we used confocal microscopy and anti-GFP/YFP immunocytochemistry. Neither mean spine density or area differed between the control group that went through the stimulation protocol (eYFP 24h-stim) and home-cage mice expressing ChR2 without light stimulus (ChR2 naive). However, chronic stimulation of ChR2-expressing mice for 24 hours (ChR2 24h-stim) significantly reduced spine density (Figure 3A; one-way ANOVA, p=0.002; Tukey’s post-hoc: eYFP 24h-stim vs. ChR2 naive, p=0.945; eYFP 24h-stim vs. ChR2 24h-stim, p=0.005; ChR2 naive vs. ChR2 24h-stim, p=0.003; n=4 mice/group); and mean spine area (Figure 3B; one-way ANOVA, p=0.009; Tukey’s post-hoc: eYFP 24h-stim vs. ChR2 naive, p=0.757; eYFP 24h-stim vs. ChR2 24h-stim, p=0.01; ChR2 naive vs. ChR2 24h-stim, p=0.03), while the proportion of different spine types did not change across groups (Figure 3C; multiple one-way ANOVAs, stubby spines, p=0.638; mushroom spines, p=0.717; thin spines p=0.465; when applying Bonferroni correction, p=1.0 for all three comparisons).

**Figure 3:**
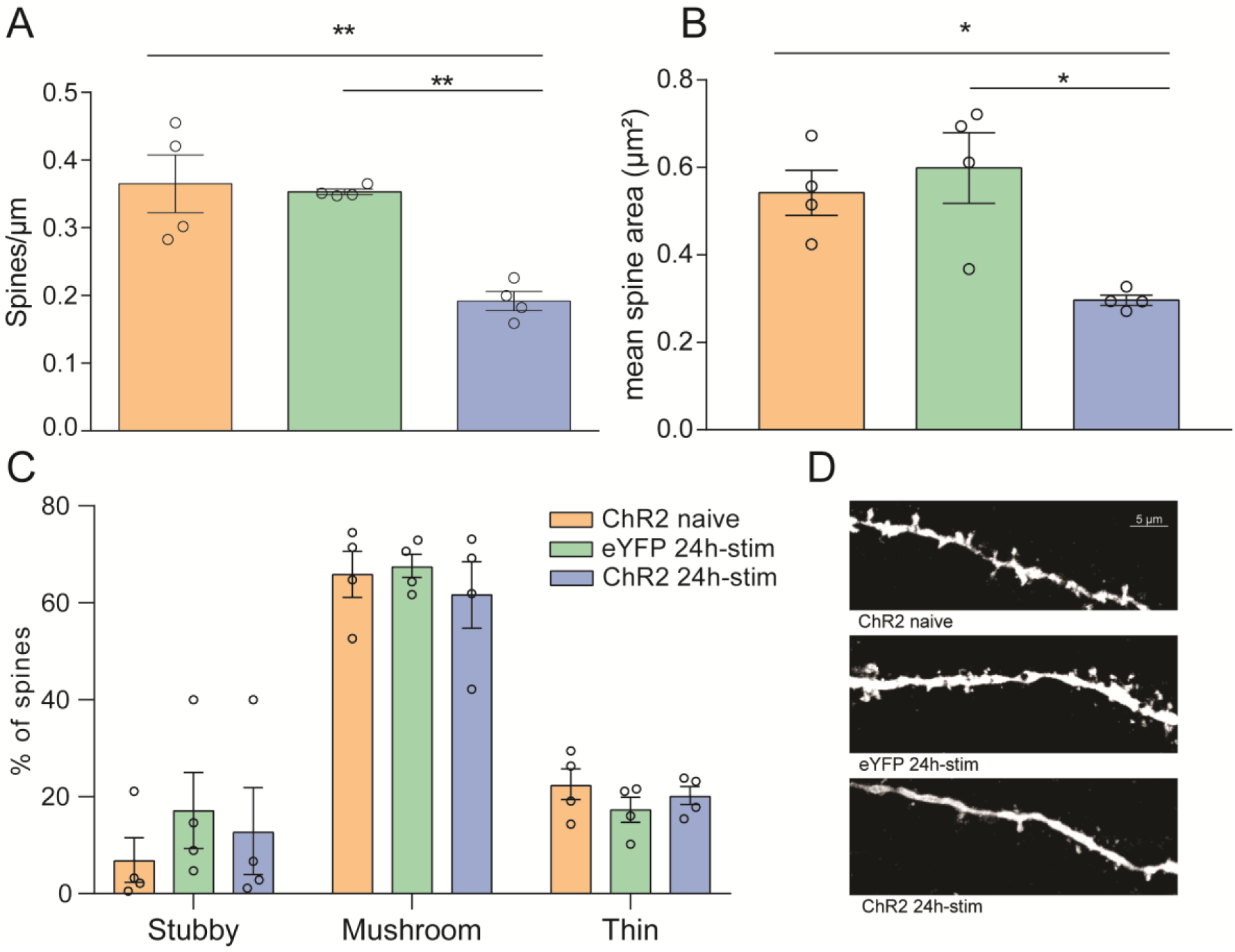
Reduced dendritic spine density and area after chronic optogenetic excitation. Three weeks after injection of viral constructs containing ChR2-eYFP or eYFP alone, mice were optogenetically stimulated for 24h (eYFP 24h-stim and eYFP 24h-stim groups). After this period, they were perfused and CA1 dendritic spines were analyzed after anti-GFP/YFP immunohistochemistry. To control for the effect of viral expression, a group of ChR2-expressing mice that did not undergo optogenetic stimulation (ChR2 naive) was included. (A) Dendritic spine density for the 3 groups. Bars show mean (± s.e.m) dendritic spine density, while circles show results for individual animals (n = 4 animals/group; 1-way ANOVA, p=0.002, Tukey’s post-hoc test; p=0.005 and p=0.003 for ChR2-naive and eYFP 24h-stim vs. ChR2 24h-stim, respectively); (B) Mean spine area (1-way ANOVA, p=0.009, Tukey’s post-hoc test; p=0.03 and p=0.01 for ChR2-naive and eYFP 24h-stim vs. ChR2 24h-stim. (C) Percentage of spine types within each group (multiple 1-way ANOVAs, stubby spines, p=0.638; mushroom spines, p=0.717; thin spines p=0.465; when applying Bonferroni correction, p=1.0 for all three comparisons). (D) Panel shows illustrative examples of dendrites from each group.

Our results are comparable to previous studies using *in vitro* neuronal excitation by optogenetics (Goold & Nicoll, 2010; Mendez et al., 2018) or picrotoxin (Fiore *et al.*, 2014), which showed reduced spine density and volume in hippocampal cultures in response to homeostatic regulation. Taken together, these morphological changes on dendritic spines may represent a fundamental mechanism of synaptic downscaling. By preserving subtype distribution, shrinking spine volume, and reducing spine density, possibly by arresting new spine growth (Mendez et al., 2018), a neuron is able to maintain and scale ongoing synaptic processes while preventing new connections, ultimately reducing its firing rate back to usual conditions.

To evaluate changes in synaptic transmission, we recorded miniature excitatory post-synaptic currents (mEPSCs) from CA1 neurons expressing either ChR2-eYFP or eYFP alone in *ex vivo* hippocampal slices obtained after the 24-h *in vivo* stimulation protocol. We observed that the amplitude for both mixed mEPSCs (containing AMPAR and NMDAR components) and AMPAR-only mEPSCs (measured after addition of the NMDAR antagonist APV) were significantly reduced in neurons from chronically stimulated animals (Figure 4A; two-way ANOVA, p= 0.0018; Tukey’s post-hoc: p=0.016 and p=0.039, respectively; n= 4 and 6 mice/group). Ratios between mixed and AMPA-only mEPSCs were higher in the ChR2 group, also indicating a reduction of the NMDAR current (Figure 4B; Student’s t test, p= 0.0001; n= 4 and 6 mice/group). Accordingly, both components have been shown to be downregulated by 24-h, 3-Hz optogenetic excitation *in vitro* (Goold & Nicoll, 2010). Albeit sparser, our chronic optogenetic stimulation protocol seems to induce similar homeostatic changes in glutamate receptors.

**Figure 4:**
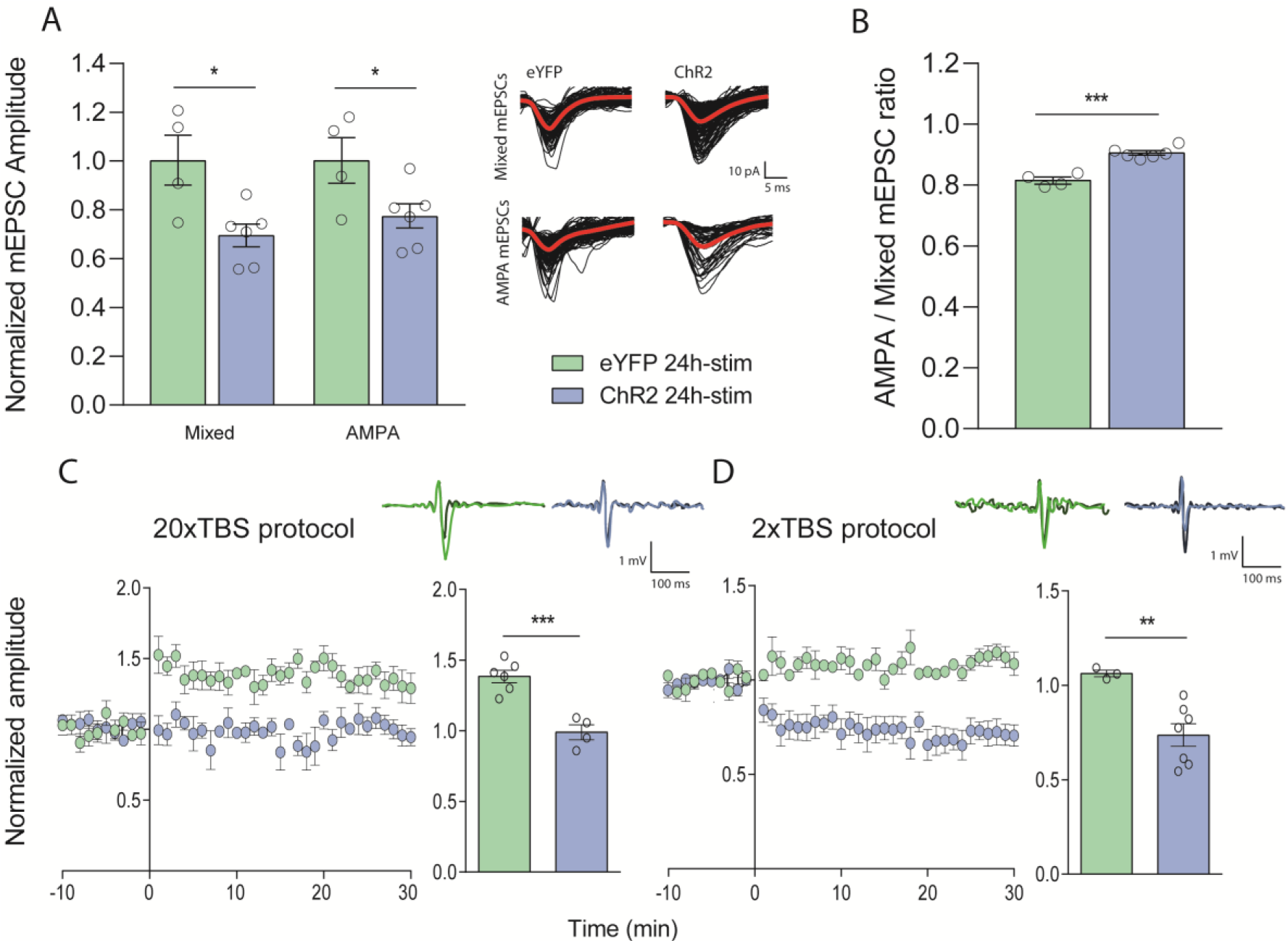
Chronic optogenetic excitation reduces mEPSC amplitude and modulates LTP and LTD. A) Amplitudes of mixed mEPSCs (containing both NMDA and AMPA components, left) and AMPA mEPSCs (right) from CA1 pyramidal cells of eYFP-control and ChR2 groups. Bars show mean (± s.e.m) amplitudes across animals, while circles show mean results for individual mice. Right panel shows representative recordings mEPSCs (black lines) and average amplitudes (red lines) for both groups. (C, D) Mean CA1 fEPSP responses in slices from eYFP-control and ChR2 animals 10 min before and 30 min after different stimulation protocols on the Schaffer collaterals. Bars show mean amplitude change from baseline (± s.e.m.) over 30 min after stimulation in each group, while circles show results for individual animals. Examples of fEPSP traces recorded before (darker lines) and after (lighter lines) the stimulation protocol are shown above the summary results. In (C) LTP, was induced in the control group after 20x-TBS, increasing field amplitude response by about 40%. In slices from mice that underwent chronic optogenetic stimulation, no change was observed (n = 4 (ChR2 24h-stim) or 6 (eYFP 24h-stim) animals; Student’s t test; p = 0.0004). In (D), after a 2x-TBS protocol, control slices did not change their fEPSP responses. However, LTD was induced in the slices of chronically stimulated mice. (n = 3 (eYFP 24h-stim) or 7 (ChR2 24h-stim) animals; Student’s t test, p = 0.0083).

Finally, to assess the effects of chronic optogenetic excitation on Hebbian plasticity, we recorded field evoked post-synaptic potentials (fEPSPs) in CA1 in response to Schaffer collateral stimulation, with baseline responses registered for 10 min using 200-ms pulses every 20 s. We then applied a 20x-theta-burst (TBS) protocol consisting of bursts of 4 pulses at 100Hz with 200-ms intervals to induce potentiation. This led to an increase in the fEPSP amplitudes in slices from the eYFP-only group, but not in those of chronically-stimulated mice (Figure 4C; Student’s t test, p = 0.0004, n = 4 (ChR2 24h-stim) or 6 (eYFP 24h-stim) animals). To investigate if chronic optogenetic activation would change the response to weak presynaptic stimulation, we used a 2x-TBS protocol that should not elicit any fEPSP amplitude increase in normal conditions. As expected, slices from chronically-stimulated eYFP animals did not show significant changes after this protocol; however, slices expressing ChR2 showed a surprising depotentiation in fEPSP responses (Figure 4D; Student’s t test, p = 0.0083, n= 3 (eYFP 24h-stim) or 7 (ChR2 24h-stim) animals).

Our results indicate that chronic *in vivo* optogenetic stimulation can drive homeostatic changes in hippocampal neurons and modulate the induction of long-term synaptic changes. To our knowledge, this is the first evidence of a direct effect of chronic neuronal stimulation on Hebbian plasticity in the hippocampal structure. The effects of chronic excitation in disturbing LTP and facilitating LTD are consistent with the hypothesis that is associated with changes in the thresholds for potentiation and depression of neuronal connections (Keck et al., 2017). This topic has been reasonably explored on sensory deprivation studies, in which recent evidence suggest that multiple processes for homeostatic regulation of excitatory synaptic strength coexist (Li *et al.*, 2014; Bridi *et al.*, 2018). On the other hand, little is known about the effect of chronic excitation on Hebbian plasticity, as it is not straightforward to induce overstimulation of sensory systems. Our study provides additional evidence to the list of commonalities between the sliding threshold model and the literature on homeostatic synaptic scaling.

Further investigation is required to describe the molecular mechanisms, circuitry modifications and behavioral consequences of this metaplastic effect, as well as to investigate if it is caused by synaptic scaling phenomena as described *in vitro* or involves Hebbian processes as well (Bridi et al., 2018). Moreover, although we have detected changes in excitatory synapses, it is also possible that circuit alterations involve changes in inhibition (Marder & Buonomano, 2004; Keck *et al.*, 2017). Previous reports of synaptic downscaling after optogenetic stimulation have conflicting indications on whether inhibitory/excitatory balance is preserved (Barral & Reyes, 2016) or altered (Mendez *et al.*, 2018), and studying these aspects our model could help in solving these controversies.

In terms of molecular mechanisms, various receptors and pathways have been involved in the modulation of both synaptic scaling and Hebbian plasticity, such as voltage-gated calcium channels (Frank, 2014), calmodulin-dependent kinases/phosphatases (Sanderson *et al.*, 2018) and others (Siddoway *et al.*, 2014). Moreover, although *in vitro* synaptic scaling has been described as NMDA receptor-independent (Turrigiano et al., 1998), recent evidence suggests that *in vivo* homeostatic plasticity caused by visual deprivation can involve NMDA-dependent mechanisms (Bridi et al., 2018). These molecules may provide interesting opportunities for pharmacological experiments to more clearly delineate whether the effects of *in vivo* chronic stimulation to occur are mostly due to synaptic scaling or Hebbian-like mechanisms.

Finally, whether similar effects also occur in physiological and/or pathological conditions remains an open question. Fundamental processes in the healthy hippocampus, such as sleep-dependent memory consolidation, can induce long-lasting enhancements the firing rate of CA1 cells (Ognjanovski *et al.*, 2014). Chronic increases in neuronal activity can also influence several phenomena, such as learning and memory (Mendez et al., 2018), neuropathic pain processing (Xiong et al., 2017) and the progression of Alzheimer’s disease (Yamamoto et al., 2015). Thus, although optogenetic excitation is a non-physiological approach, it can be a useful tool to make homeostatic neuronal regulation – a process up to now mostly studied in cultured neurons – more amenable for *in vivo* studies that can investigate its relations with physiological processes in the brain.

## Detailed Methods

### Animals and optogenetic stimulation

Experiments were performed on C57BL/6J mice of both sexes, aged 6-10 weeks, provided by the Transgenic Animal Lab (LAT), Federal University of Rio de Janeiro, Brazil and by the Brain Institute, Natal, Brazil. All procedures were approved by the Institutional Ethics Committee for the Care and Use of Laboratory Animals of the Federal University of Rio de Janeiro (CEUA-CCS 024/15) and the Federal University of Rio Grande do Norte (Protocol 015.004/2017).

For stereotactic surgery, animals were anesthetized with i.p. injections of ketamine (100 mg/kg) and xylazine (20 mg/kg). Viral solution (1 × ?10^12^ particles/ml) was delivered unilaterally at 0.2 µl/min in the CA1 region of the ventral hippocampus (A.P. 2.8 mm; M.L. 2.6 mm; D.V. 3.0 and 3.5 mm; 0.75 µl/site) before the fiber-optic cannula was fixed. Vectors containing ChR2 under the promoter of the CaMKII-α gene (rAAV5/CaMKIIa-ChR2(H134R)-EYFP) or a control vector lacking the ChR2 sequence (rAAV5/CaMKIIa-EYFP) were obtained from the Vector Core Facility at the University of North Carolina.

After a period of 3 to 5 weeks after surgery, we applied a chronic stimulation protocol using 15-ms pulses of blue light (473 nm) every 90 s during 24 hours on freely-moving mice. Immediately after this procedure, animals were euthanized as described below.

### Dendritic spine morphology analysis

For analysis of dendritic spines, animals were anesthetized with ketamine (0.5-1g/kg) and perfused with 4% paraformaldehyde in 0.1M PBS. We obtained 150-µm-thick horizontal hippocampal slices using a vibratome, and immunocytochemistry was performed on free-floating sections with sheep anti-GFP/YFP (1:700; AbD Serotec) and Alexa Fluor 488-conjugated donkey anti-sheep IgG (1:500; Thermo-Fischer). Labeled dendrites were imaged using a Leica TCS SPE confocal microscope with a 63x-objective, followed by 1.5-2.5x digital zoom. Between 3 and 5 different fields focusing on pyramidal apical dendrites of CA1 were digitally captured for each animal.

A second experimenter, blinded to the experimental groups, chose at least three images per animal based on their quality for subsequent analysis. Due to the high-density expression of labeled dendrites, it was not possible to individually identify the originating cells; thus, mean spine density was calculated for each field by dividing the spine number of all clearly visible dendrites by their total length. To measure mean spine area in each field, images were binarized and spines were isolated as described in Hsia *et al.* (1999). The average of all fields was used to compute mean spine density and area for each mouse. For the estimation of dendritic spine types, we classified spines as mushroom, thin or stubby spines as described in Nimchinsky *et al.* (2002), calculating the percentage of clearly-defined spine types for each animal. All assessments were made using ImageJ by an observer who was also blinded to the experimental groups.

### Electrophysiology

For electrophysiology, mice were euthanized by ketamine (0.5-1g/kg) and then decapitated. Horizontal 400-µm-thick hippocampal slices were obtained with a vibratome (Leica Biosystems) and maintained in artificial cerebrospinal fluid (aCSF: 124mM NaCl, 3.5mM KCl, 1.25mM NaH2PO4, 1.5mM MgCl2, 1.5mM CaCl2, 24mM NaHCO3 and 10mM glucose), constantly bubbled with 95% O2 and 5% CO2.

For qualitative demonstration of ChR2-mediated inward currents, borosilicate glass electrodes (resistance = 4–8 MΩ) were filled with an internal solution containing 130mM K-gluconate, 7mM NaCl, 2mM MgCl2, 2mM ATP, 0.5mM GTP, 10mM HEPES, 0.1mM EGTA, and pH was adjusted to 7.2 using KOH. Postsynaptic currents were obtained in voltage clamp at a holding potential of -60 mV.

For field spike measurements after *ex vivo* optogenetic stimulation, we recorded the extracellular LFP by placing ACSF-filled borosilicate glass electrodes in the CA1 stratum pyramidale of slices expressing ChR2. After recording LFPs for 1 s, a 15-ms light pulse was delivered and the recording was maintained for another 1 s. After 5 trials of this protocol for each slice, the mean number of spikes before and after stimulations was registered using the automatic threshold algorithm described by Quiroga et al. (2007).

For mEPSC recordings, the same internal solution used for demonstration of ChR2 currents was used, and tetrodotoxin (1 µM) and bicuculline methochloride (10 µM) were added to the bath. YFP-expressing CA1 pyramidal cells were patched and recordings were made in voltage-clamp at holding potential of -60 mV for at least 5 min with 5x-gain and filtered at 1 kHz. To isolate AMPAR mEPSCs, NMDA receptors were then inhibited with D-AP5 (30 µM). The amplitudes of miniature EPSCs were measured using MATLAB (Mathworks) as described in Leão *et al.*, 2005).

For LTP experiments, extracellular fEPSP recordings were obtained by placing a concentric stimulation electrode at the stratum radiatum as previously described (Leão et al., 2012). A borosilicate glass pipette filled with ACSF was used to record Schaffer collateral fEPSPs at the CA1 region, 200–400µm away from the stimulation electrode. Qualitative stimulus-response curves were visualized to confirm the robustness of the fEPSP. Stimulation strength was adjusted to obtain 50–60% of the maximum fEPSP amplitude, and 10 min of recordings were used to obtain a stable fEPSP baseline, with 200ms pulses delivered every 20 s. Synaptic plasticity was induced by 2x-TBS or 20x-TBS (2 or 20 bursts of four pulses at 100 Hz spaced by 200 ms, respectively). After stimulation, recording of stimulus-response curves continued for 30 min, which were aggregated to measure the amplitude change from baseline. When more than one slice from the same animal successfully went through the same theta-burst protocol, the average amplitude change for that mouse was used.

### Statistical Analysis

Data from all experiments were analyzed in GraphPad Prism 7. Comparisons between groups were performed using Student’s t-test, paired t-test, one-way ANOVA or two-way ANOVA, followed by Tukey’s post-hoc test, with the specific test described for each result. All raw data are available as supplementary material of the preprint version of the manuscript.

## Supporting information

## Acknowledgments

This work was funded by grants and scholarships from CNPq, CAPES and FAPERJ. The authors thank Prof. Roberto Lent for providing the laboratory infrastructure for the immunohistochemistry experiments.

## Author contributions

T.C.M., O.B.A, and R.N.L. conceived and designed the experiments. T.C.M., L.L.P., D.R., J.W., R.G.M., R.V.L.C., and R.N.L. performed experiments. T.C.M., L.L.P., and R.N.L analyzed data. T.C.M. and O.B.A wrote the manuscript. All authors revised the final version of the manuscript.

## Conflict of Interest

The authors have no conflicts of interest to declare.

