## Supplementary material for "Chronic *in vivo* optogenetic stimulation modulates neuronal excitability, spine morphology and Hebbian plasticity in the mouse hippocampus": run_slice_final.m

```
clear all

y1=load('002_baseline[1-30].mat');
y2=load('002_30min[1-90].mat');

y1=struct2cell(y1);
y1=y1{1};
response1 = zeros(length(y1)/2,1);

y2=struct2cell(y2);
y2=y2{1};
response2 = zeros(length(y2)/2,1);

c=1;
for i=2:2:length(y1)
    base=mean(y1{i}(3100:3200));
    response1(c)=base-min(y1{i}(3300:3400));
    c=c+1;
end

c=1;
for i=2:2:length(y2)
    base=mean(y2{i}(3100:3200));
    response2(c)=base-min(y2{i}(3300:3400));
    c=c+1;
end

%corre��o outliers (pnts > 5*sd)
for i=6:30
    if abs(response1(i))>abs(mean(response1(i-5:i-1)))+5*std(response1(i-5:i-1))
        response1(i)=mean(response1(i-5:i-1));
    end
end
for i=6:90
    if abs(response2(i))>abs(mean(response2(i-5:i-1)))+5*std(response2(i-5:i-1))
        response2(i)=mean(response2(i-5:i-1));
    end
end

%plot
x1=[1:length(response1)];
x2=[length(response1)+1:length(response1)+length(response2)];

plot(x1,response1,'o')
hold on
plot(x2,response2,'o')
hold off
```
