## Supplementary figures and images for "Chronic *in vivo* optogenetic stimulation modulates neuronal excitability, spine morphology and Hebbian plasticity in the mouse hippocampus"

### 1.bmp

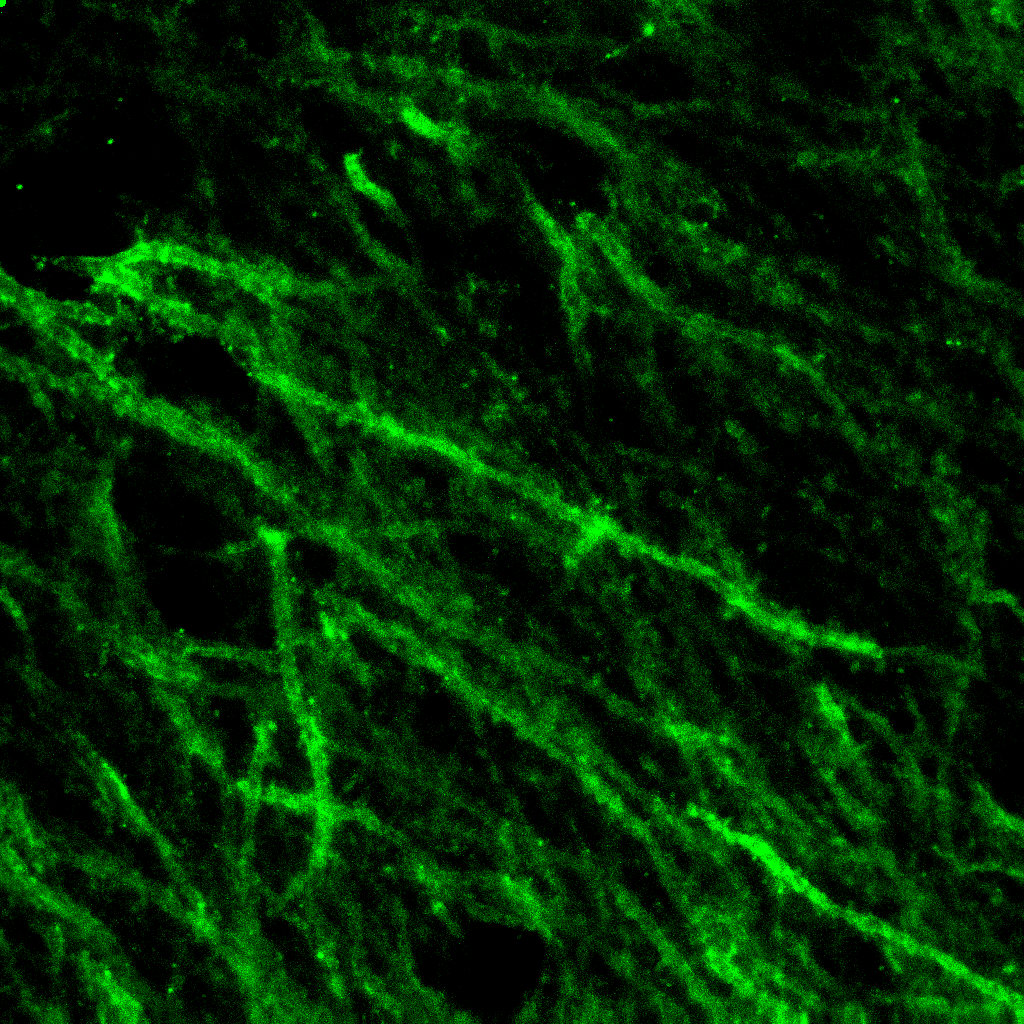

### 1.bmp

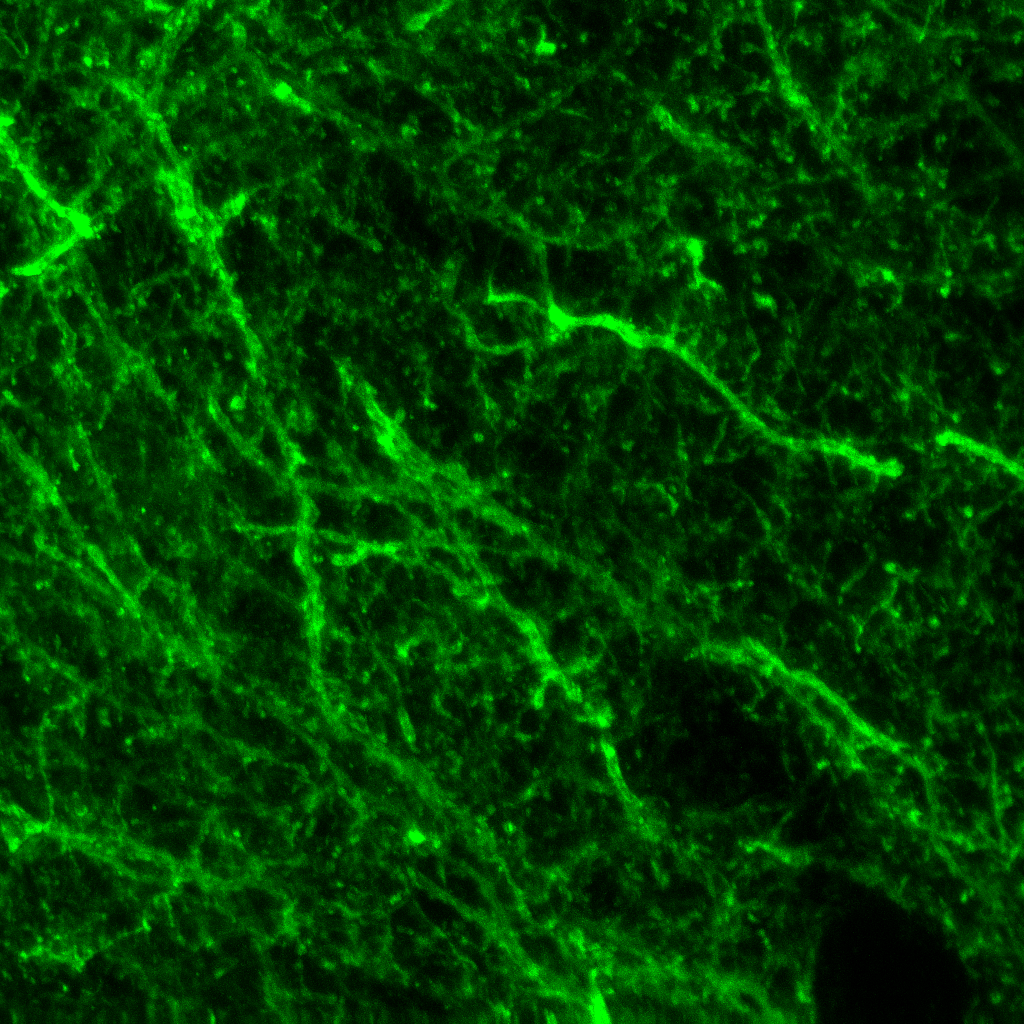

### 1.bmp

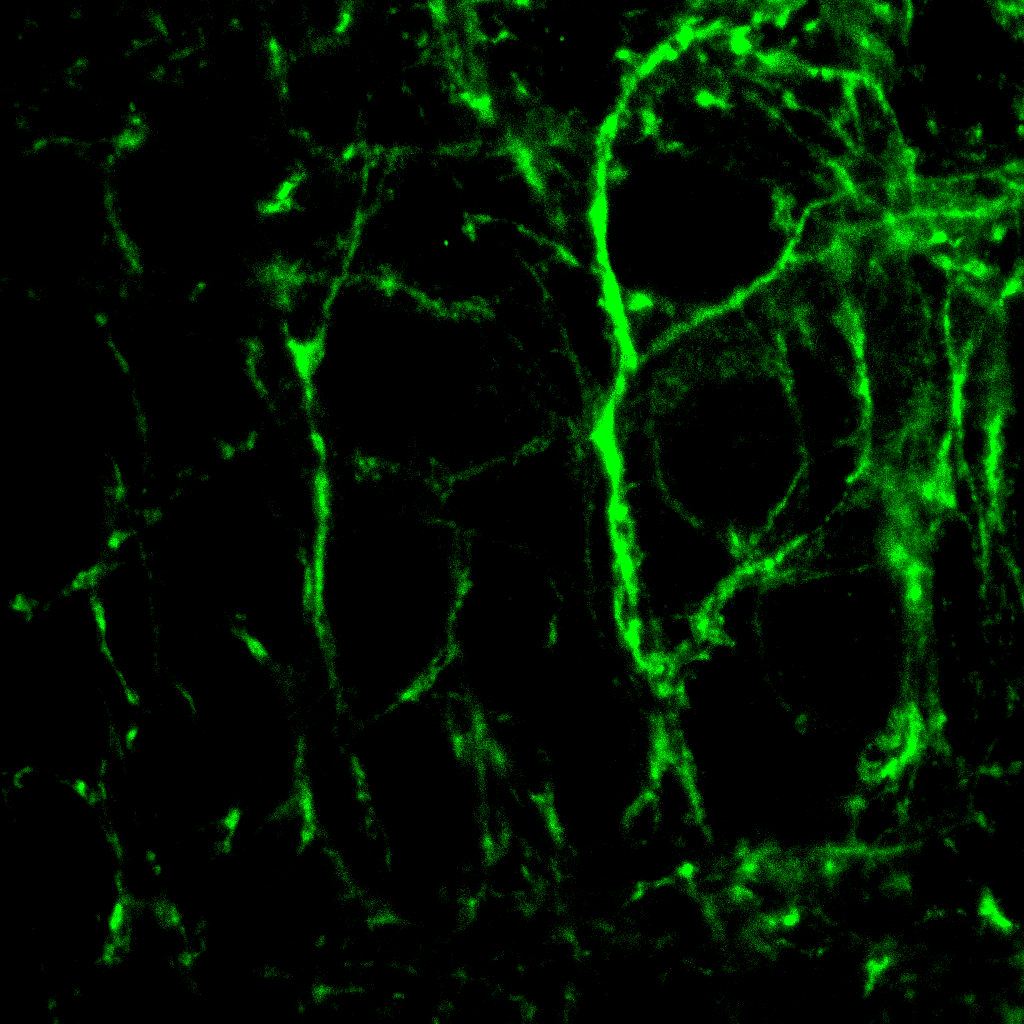

### 1.bmp

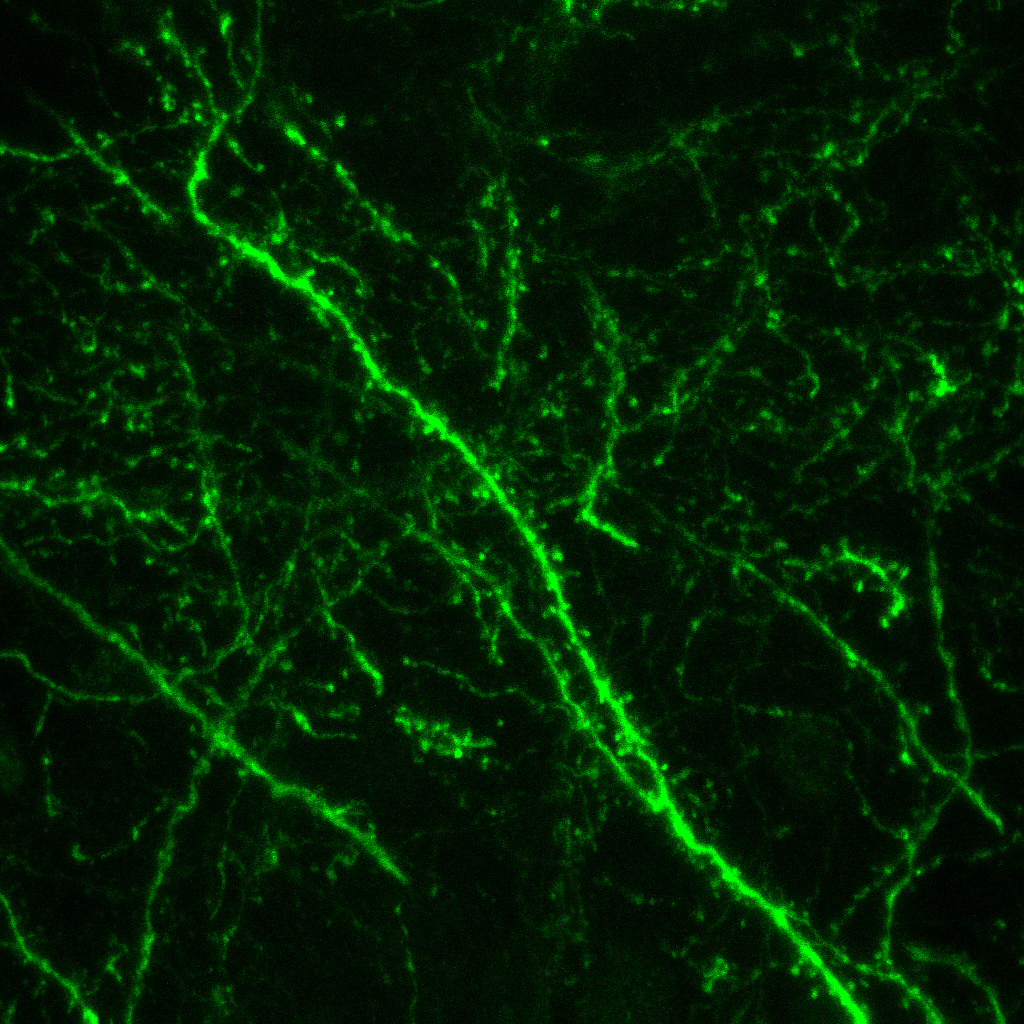

### 1.bmp

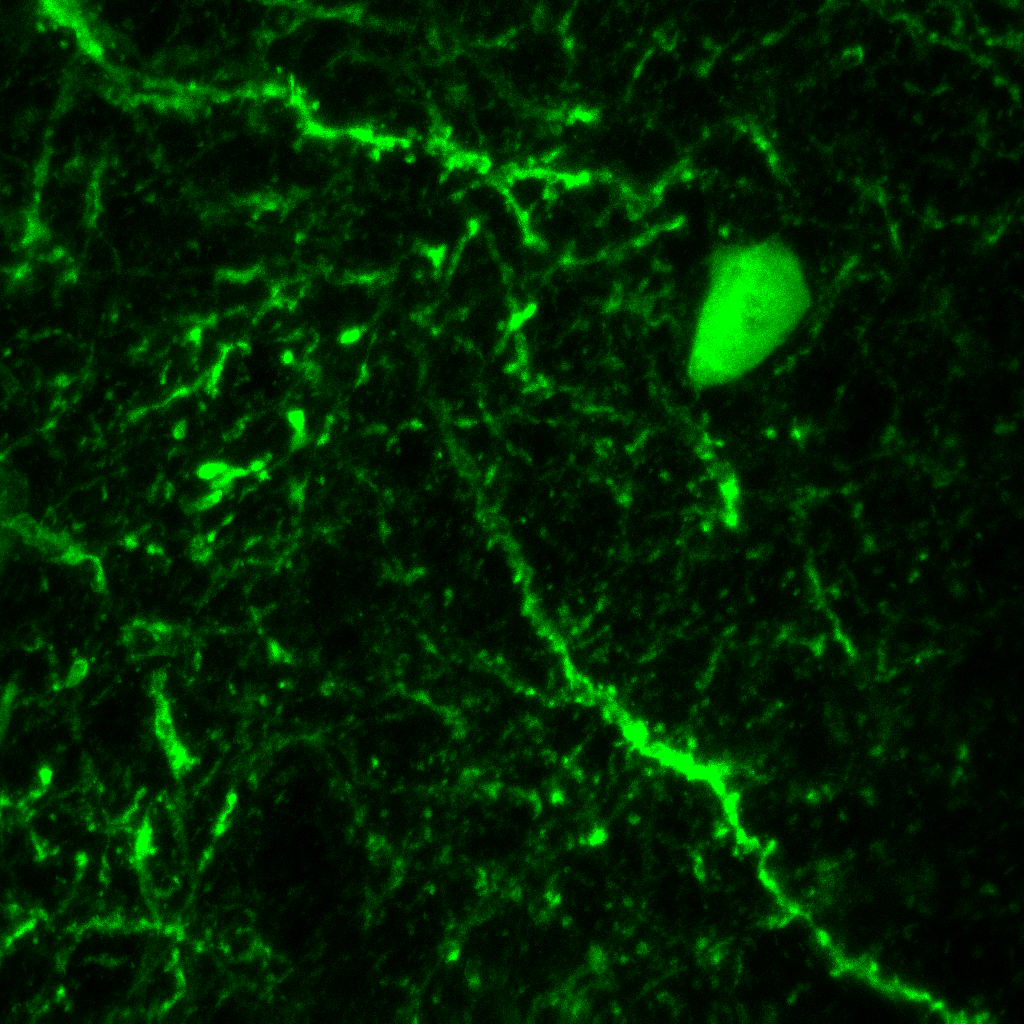

### 1.jpg

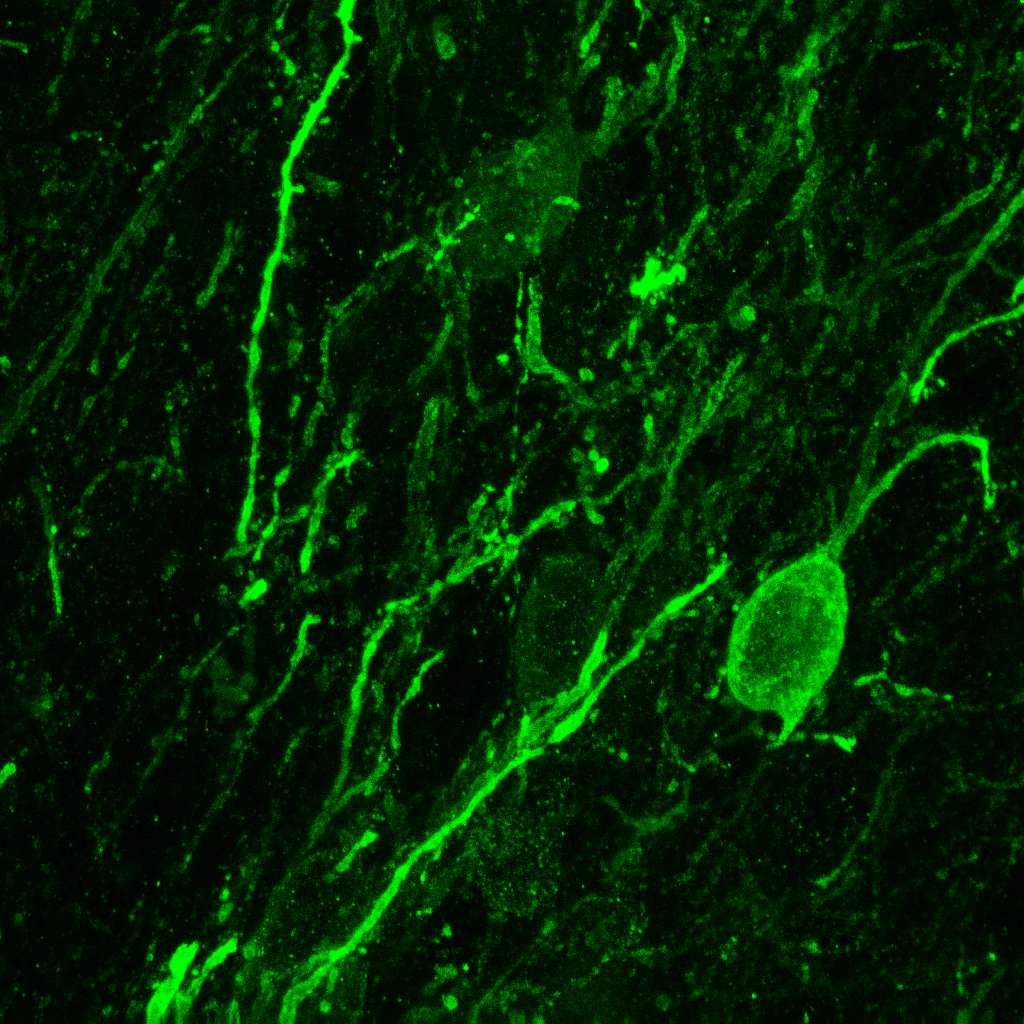

### 1.jpg

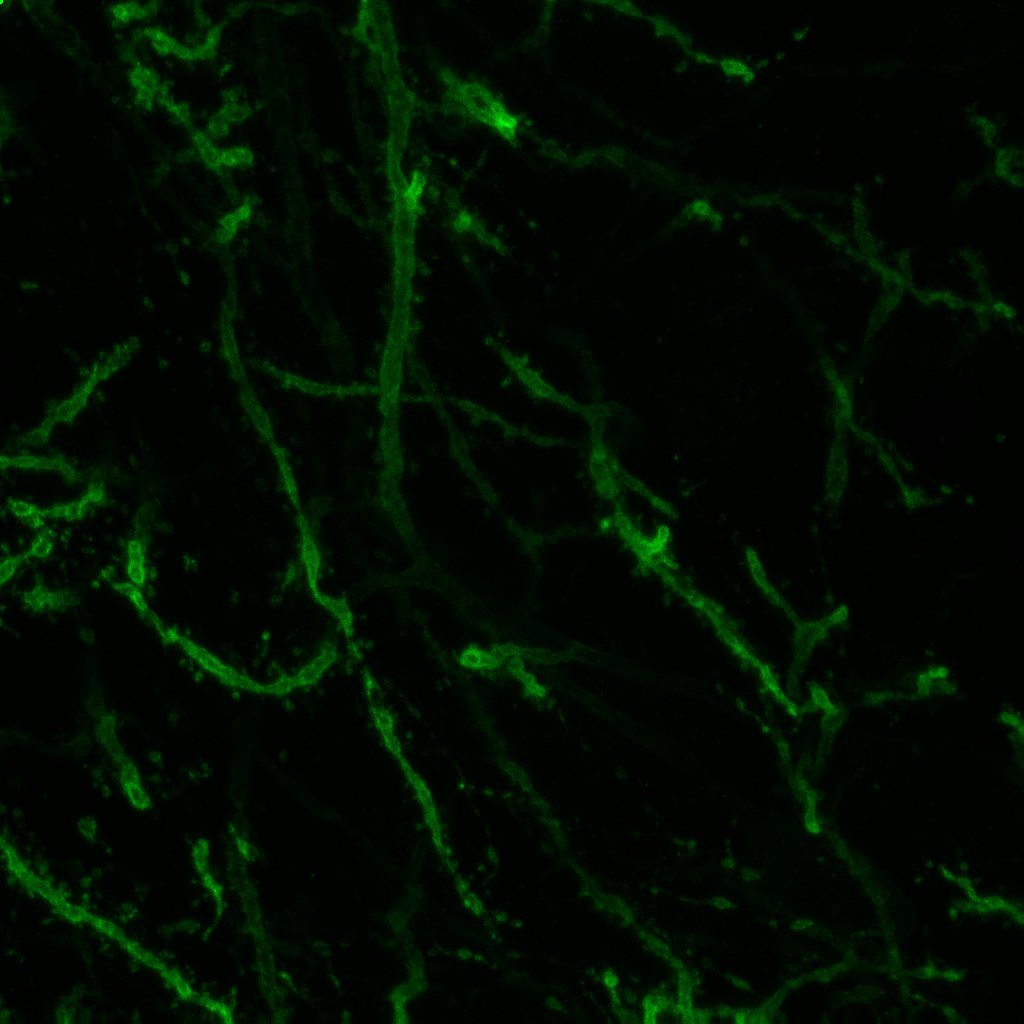

### 1.jpg

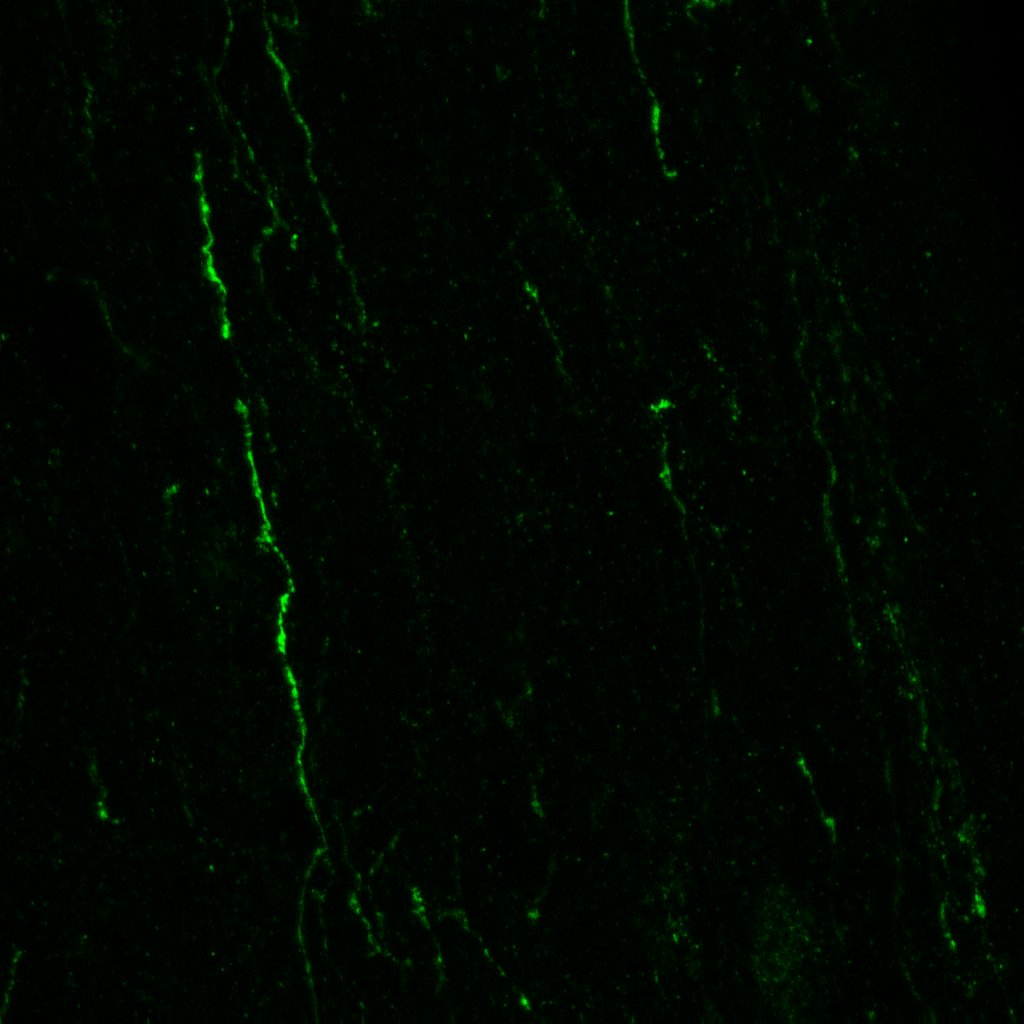

### 1.jpg

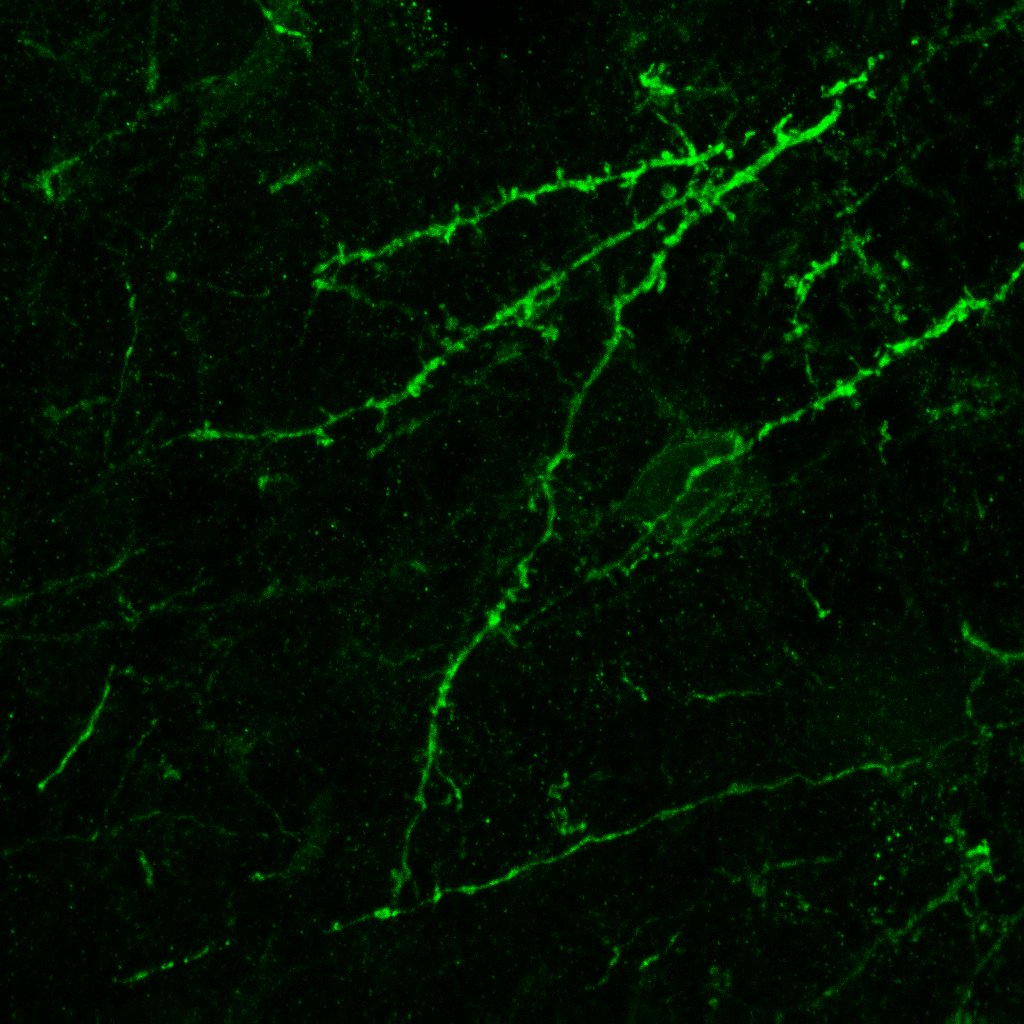

### 1.jpg

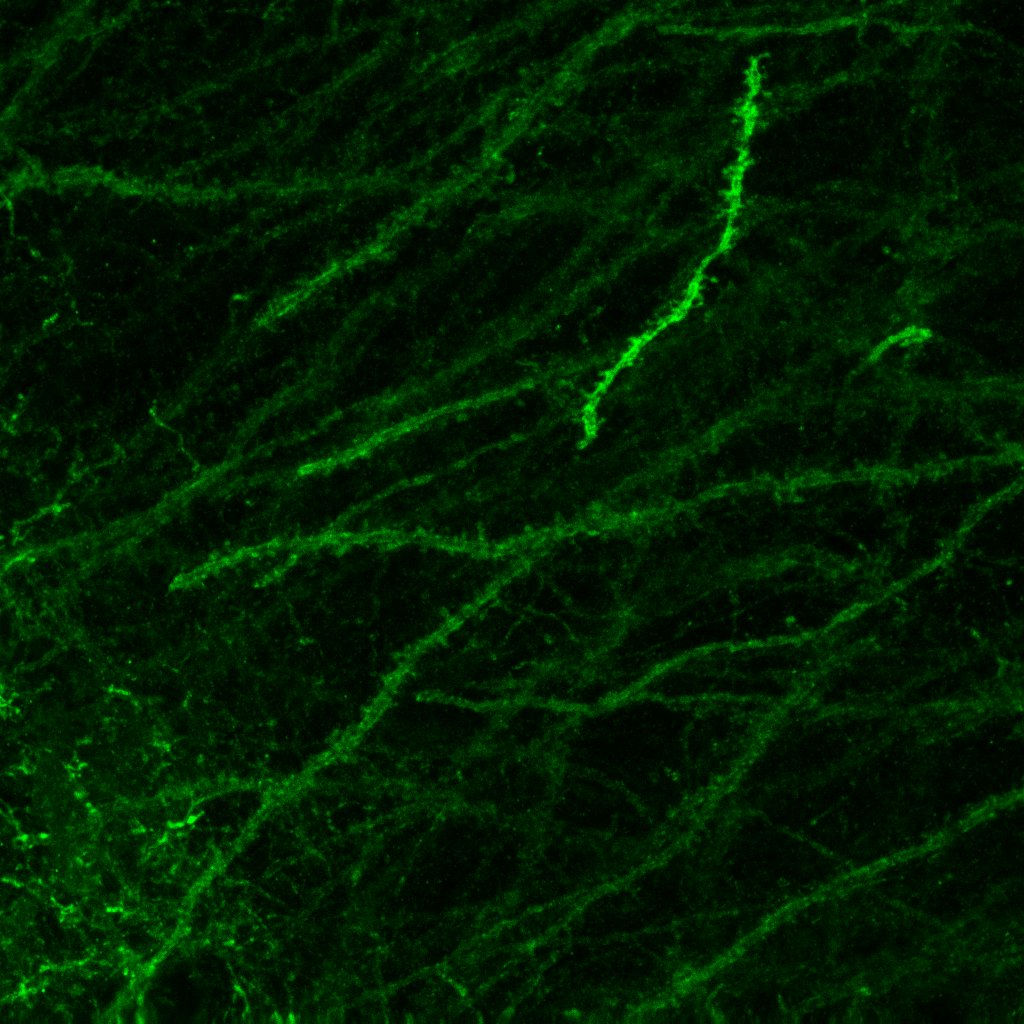

### 1.jpg

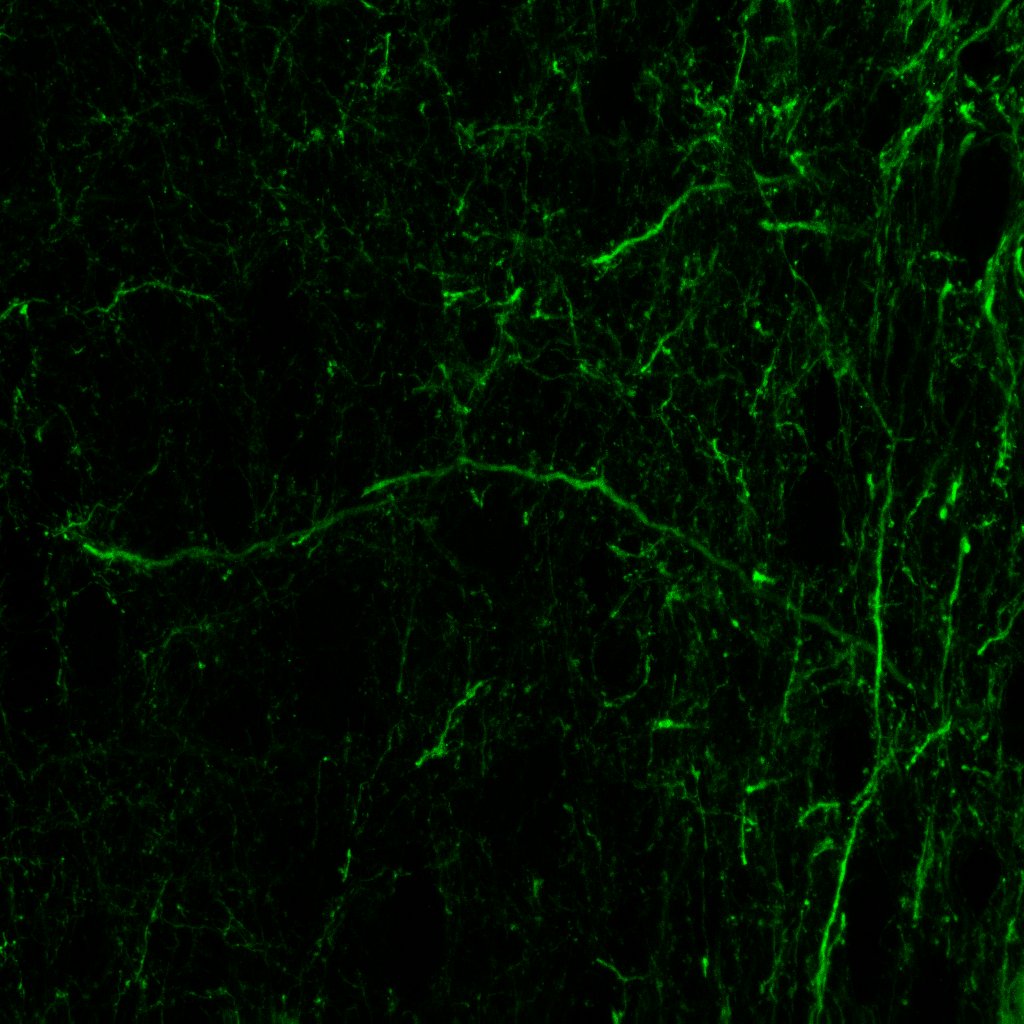

### 1.jpg

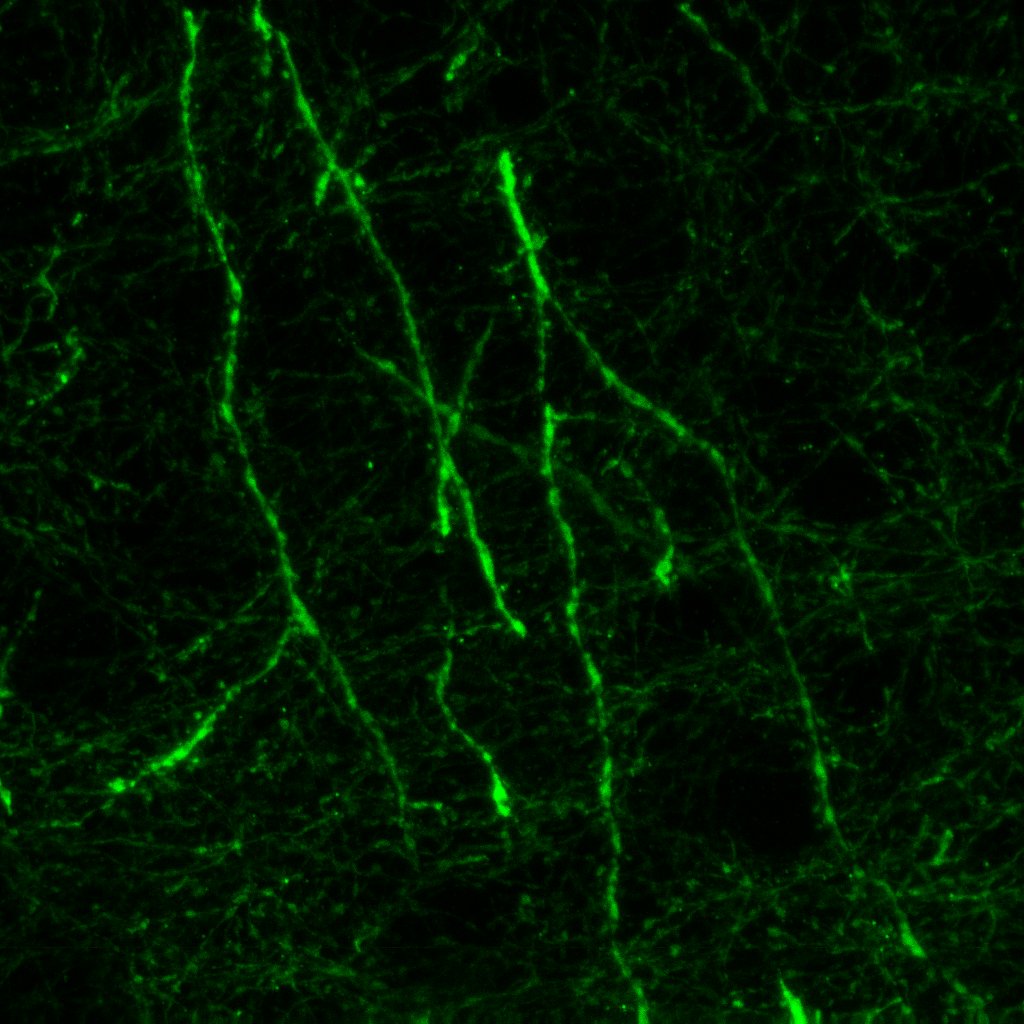

### 2.bmp

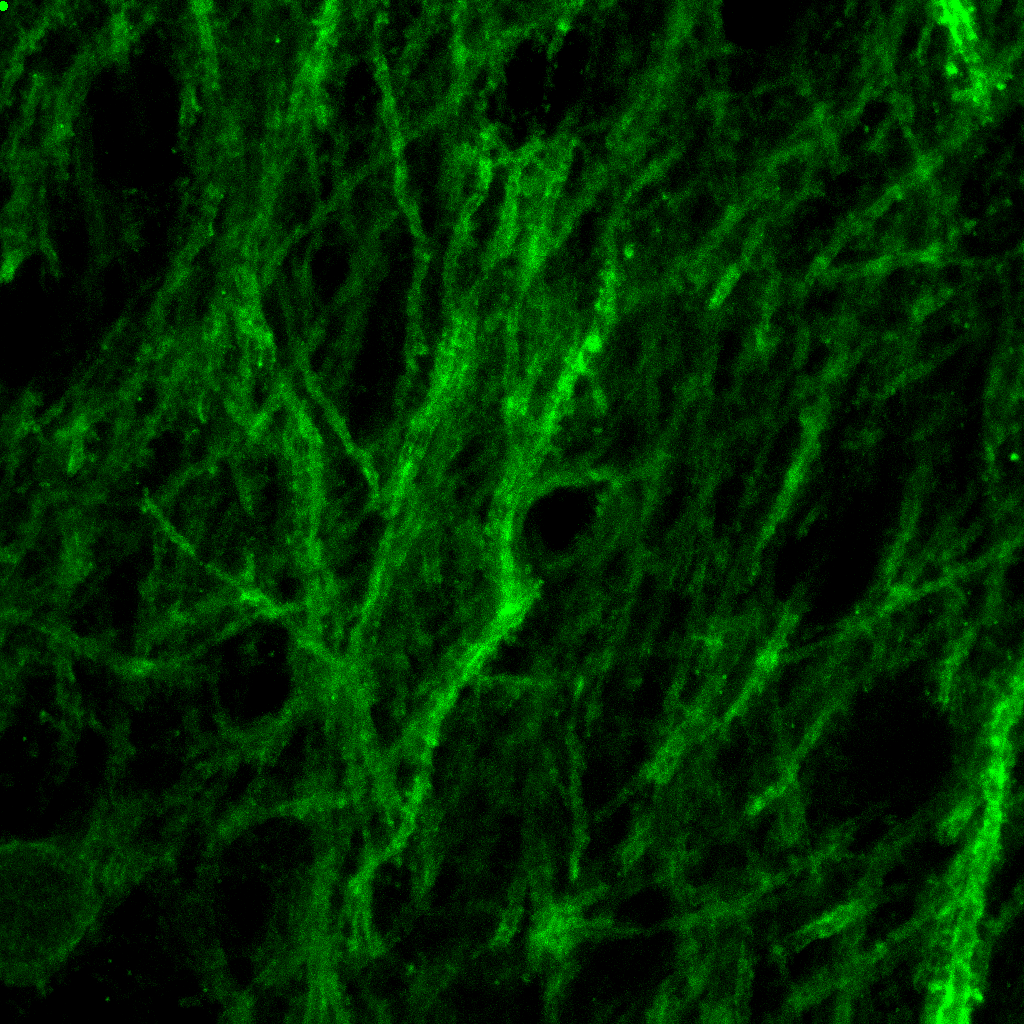

### 2.bmp

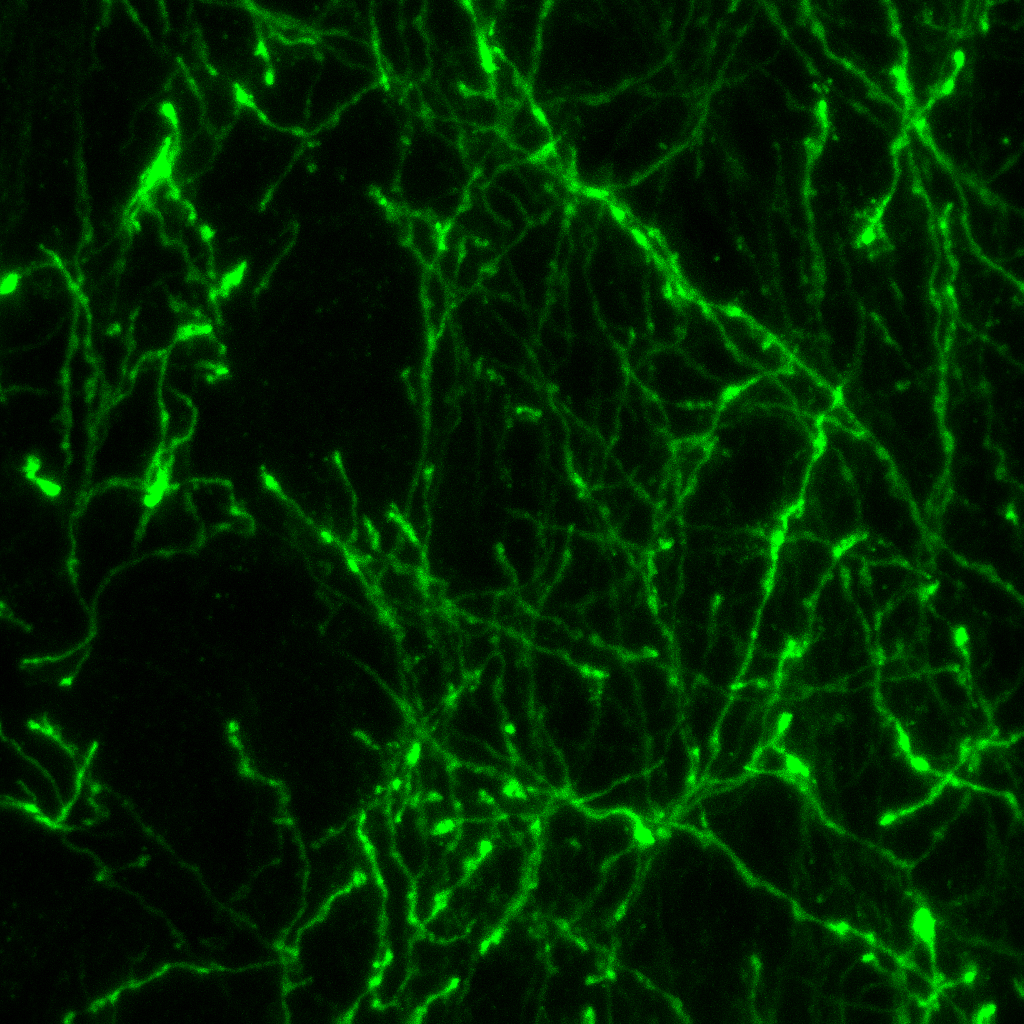

### 2.bmp

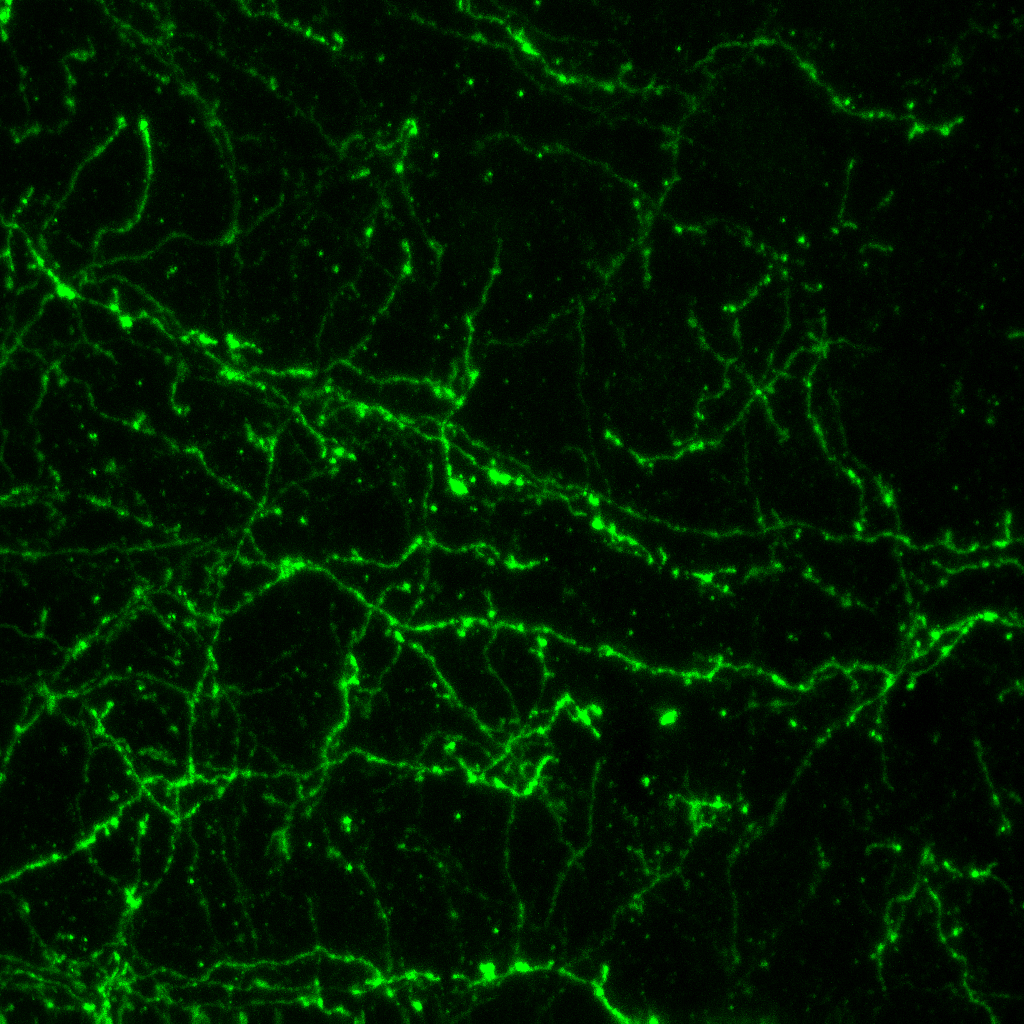

### 2.bmp

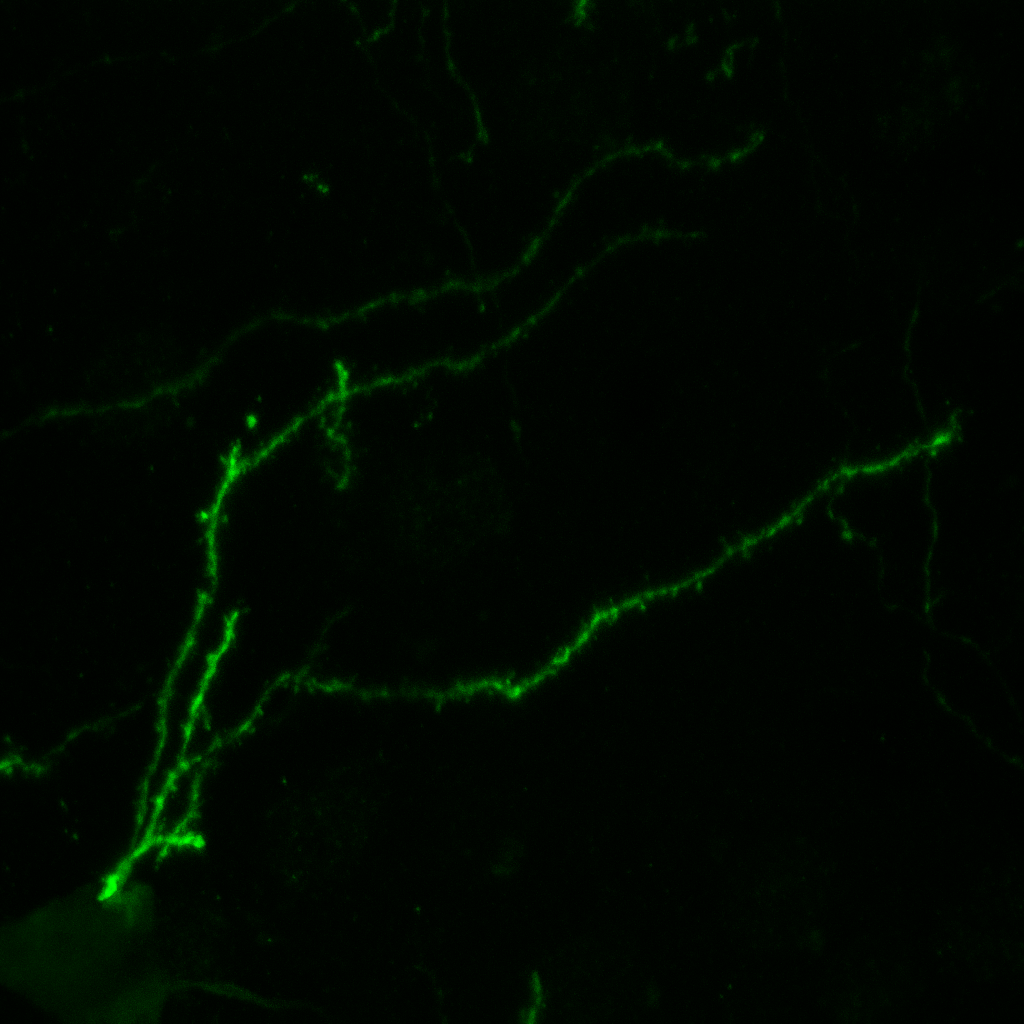

### 2.bmp

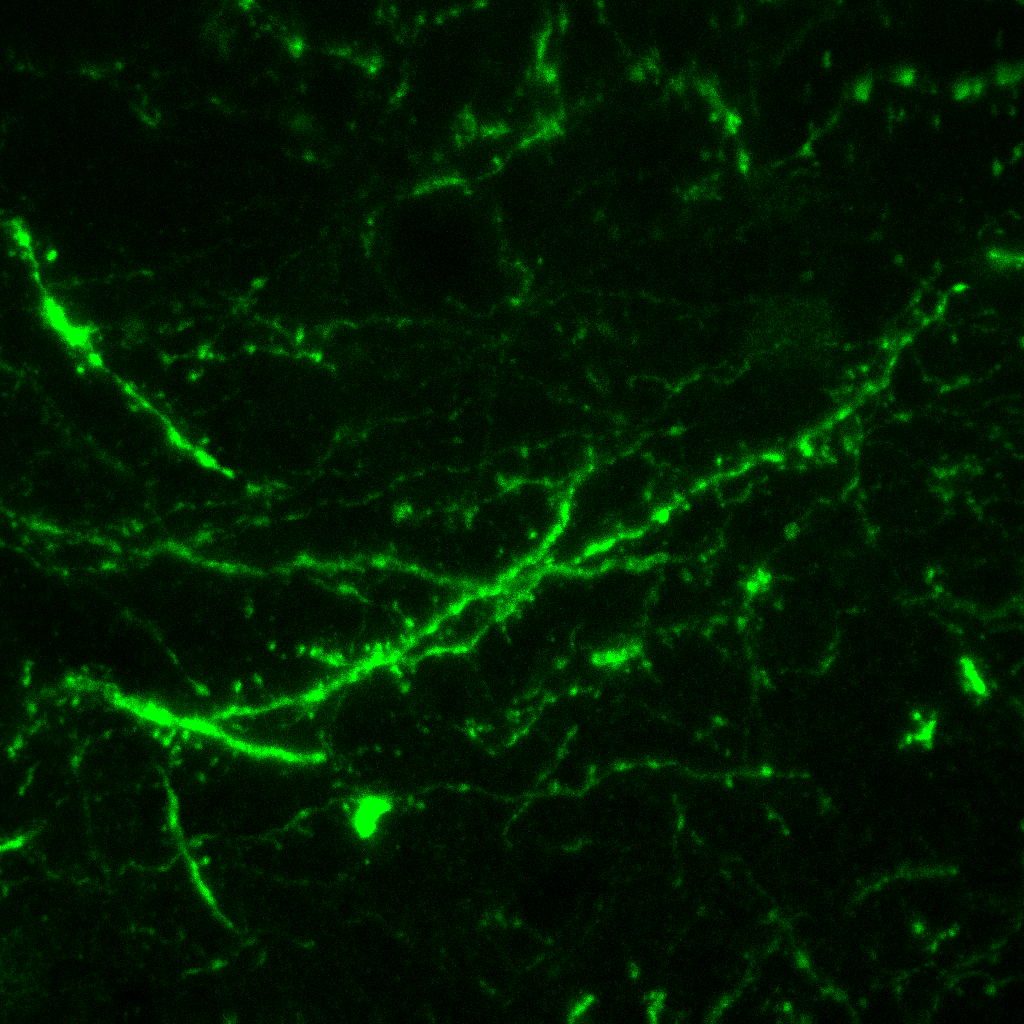

### 2.jpg

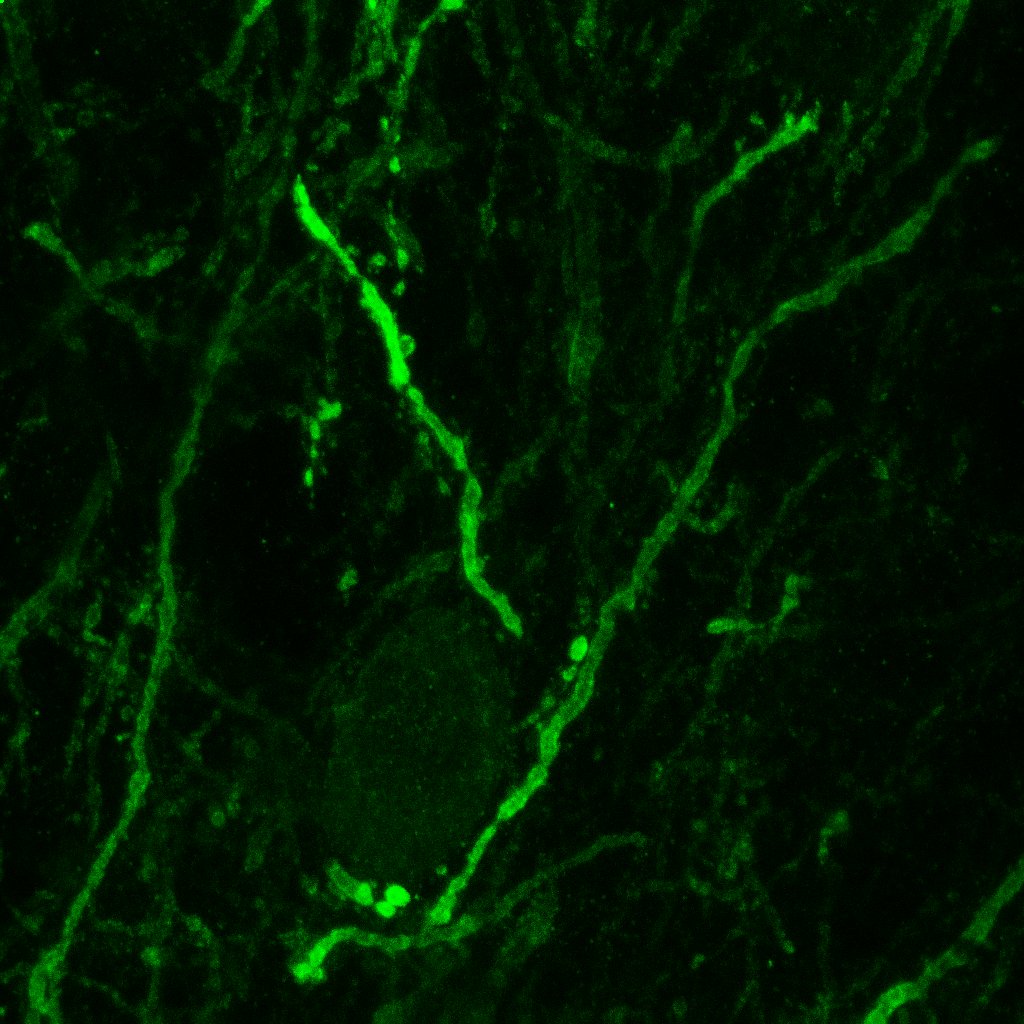

### 2.jpg

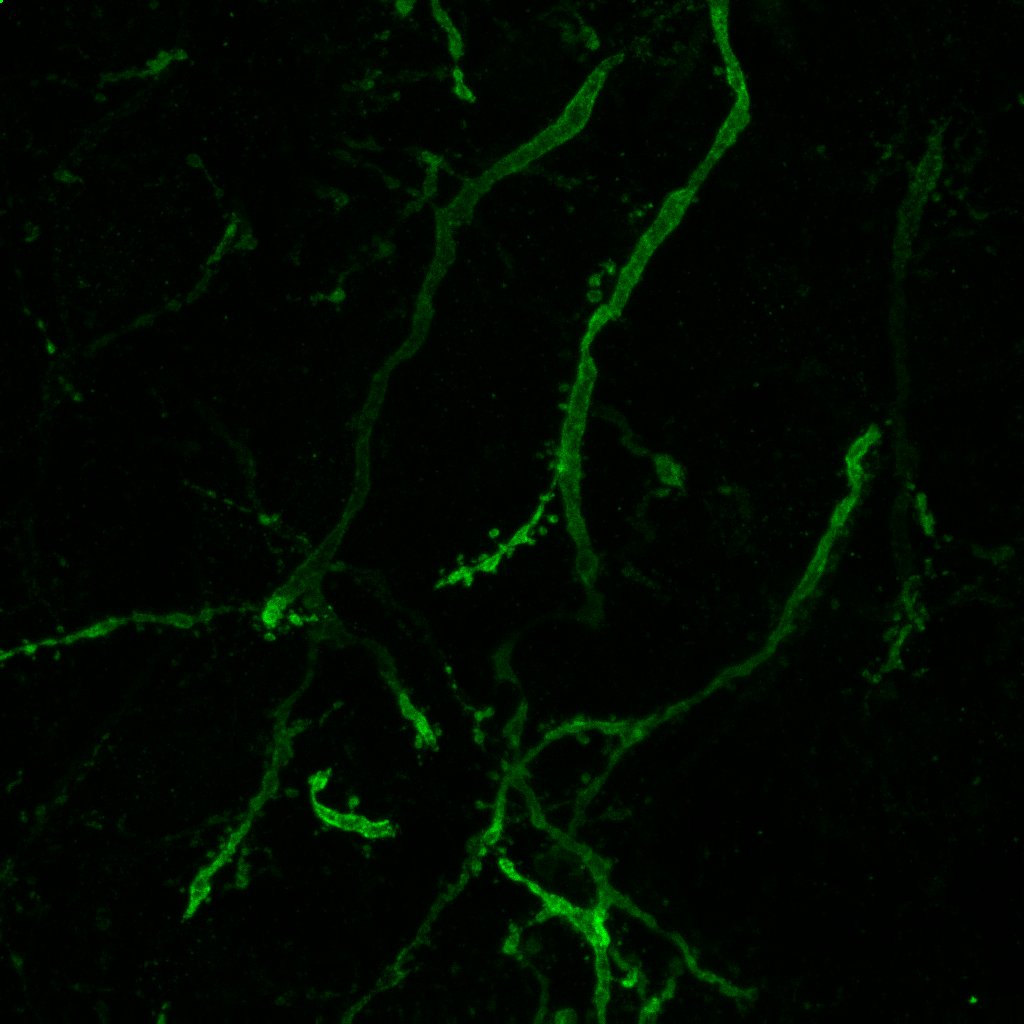

### 2.jpg

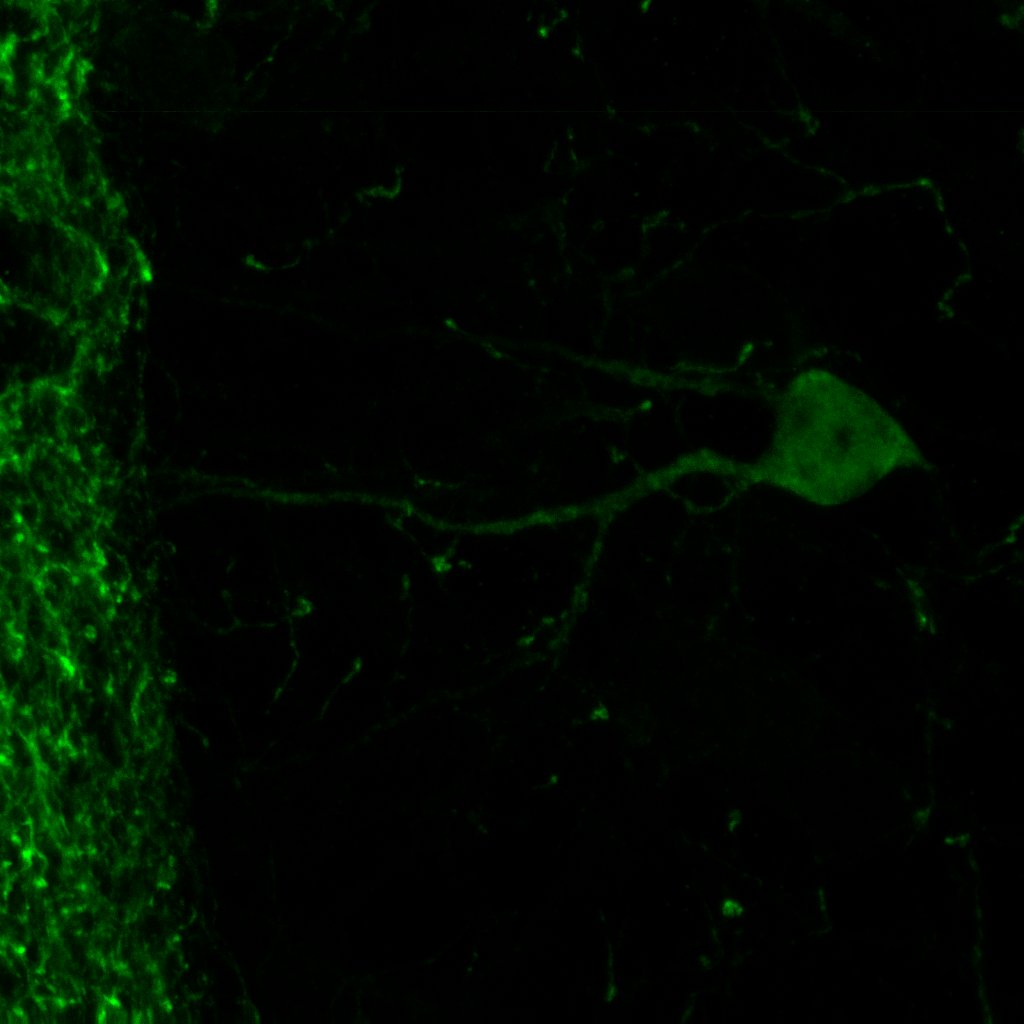

### 2.jpg

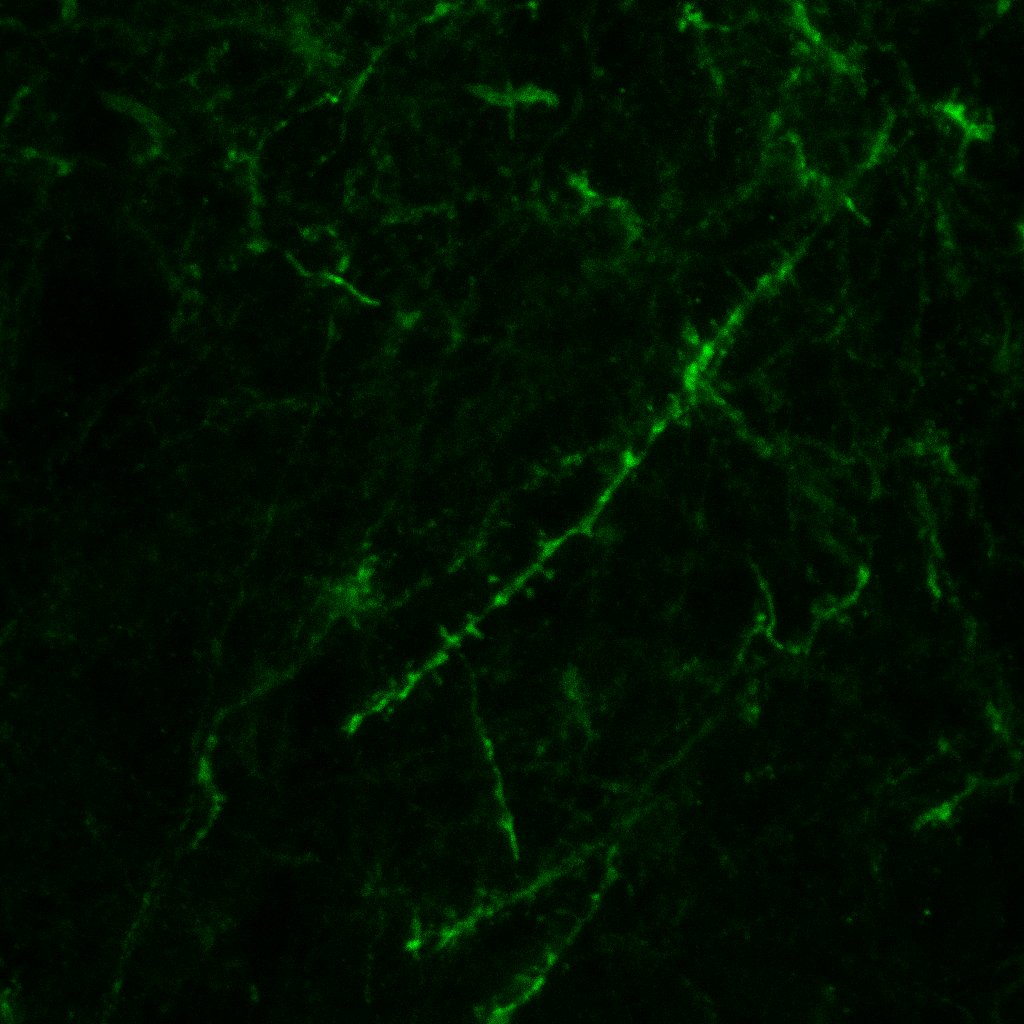

### 2.jpg

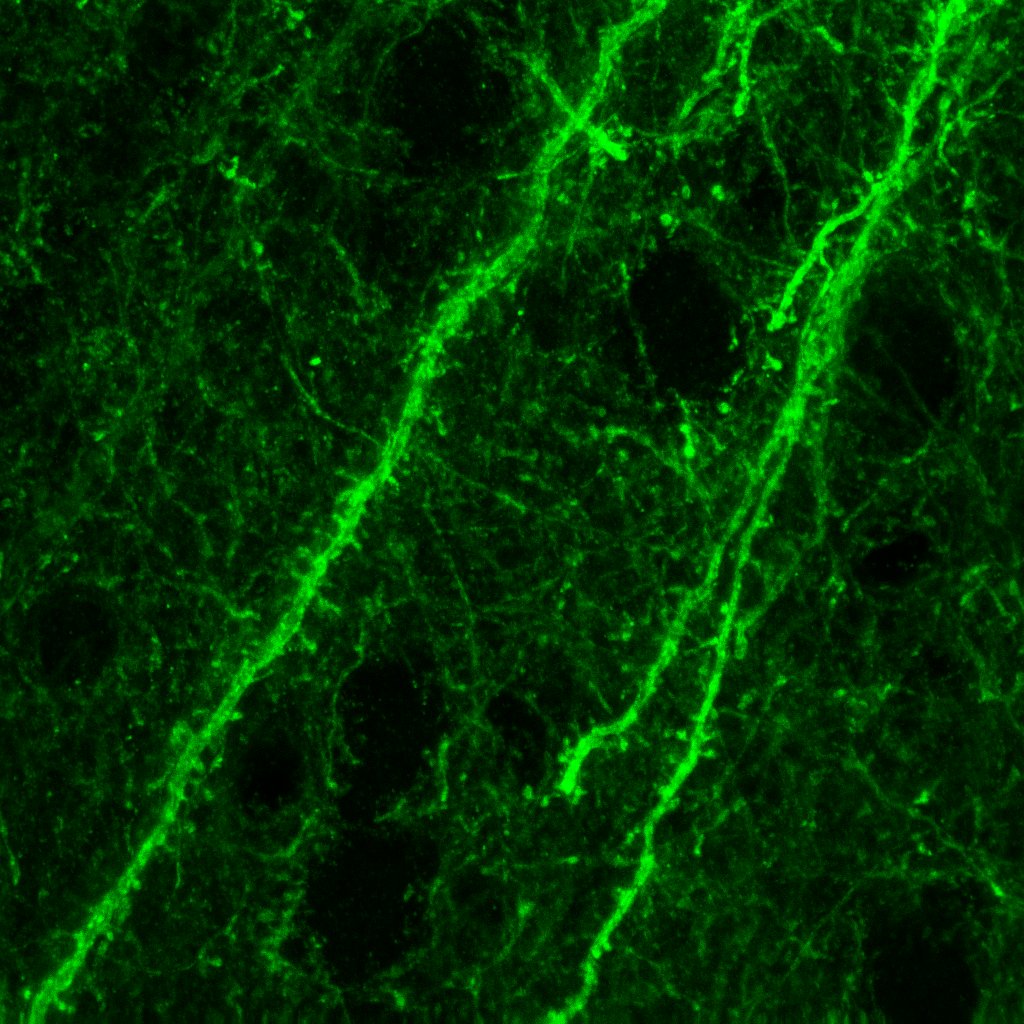

### 2.jpg

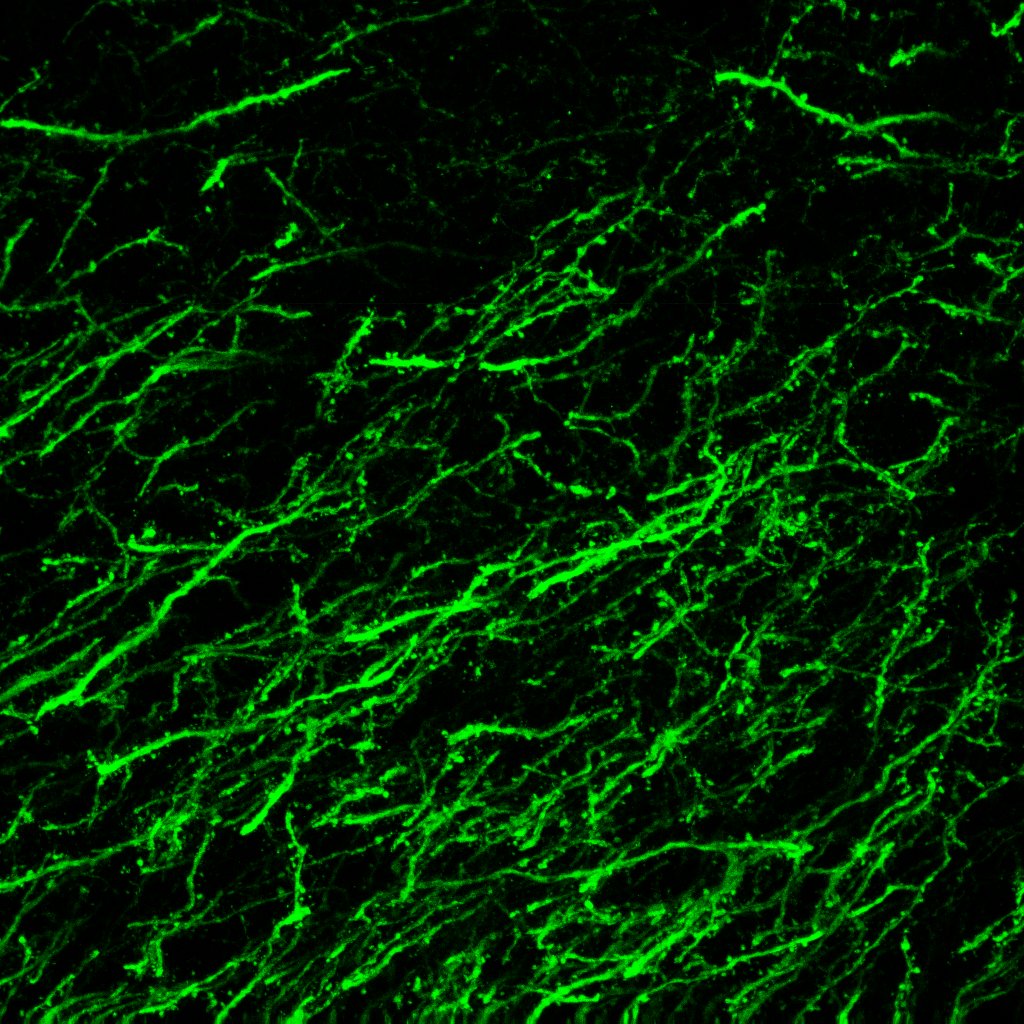

### 2.jpg

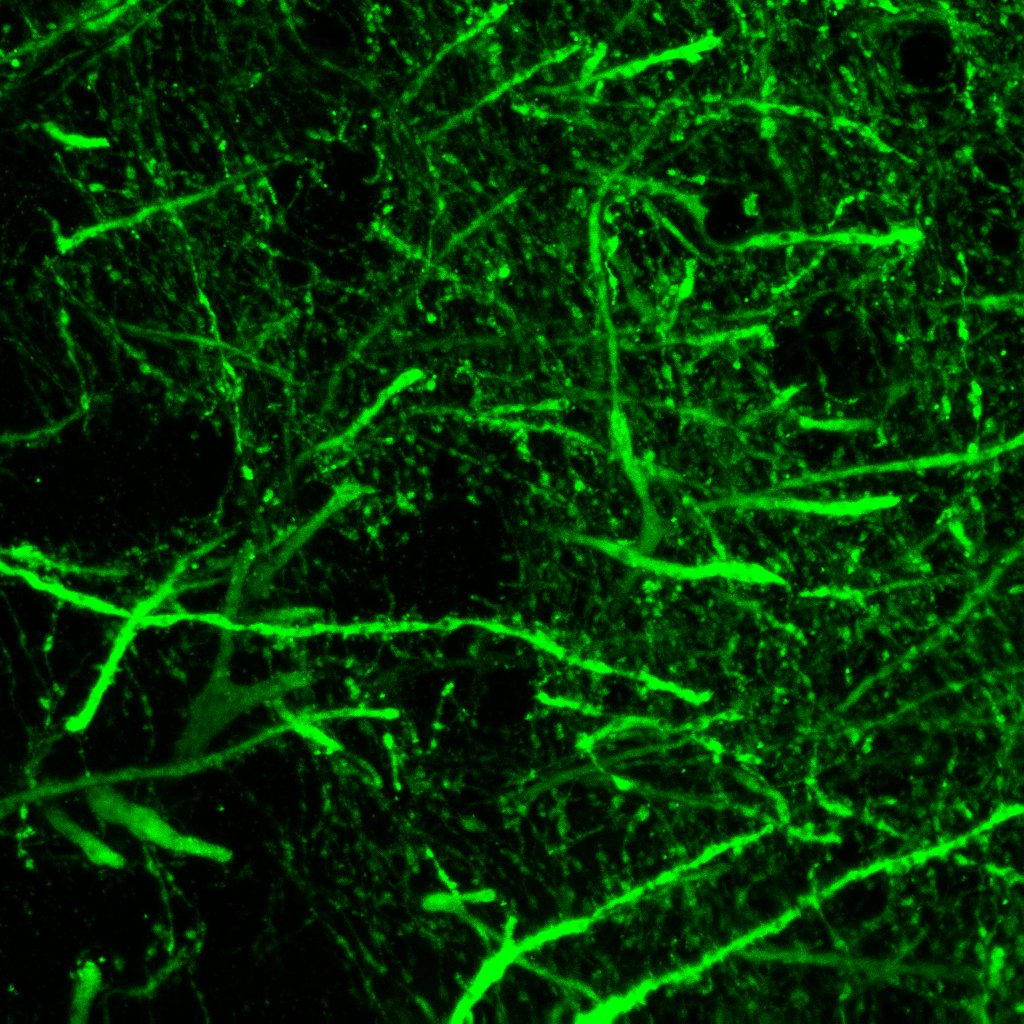

### 3.bmp

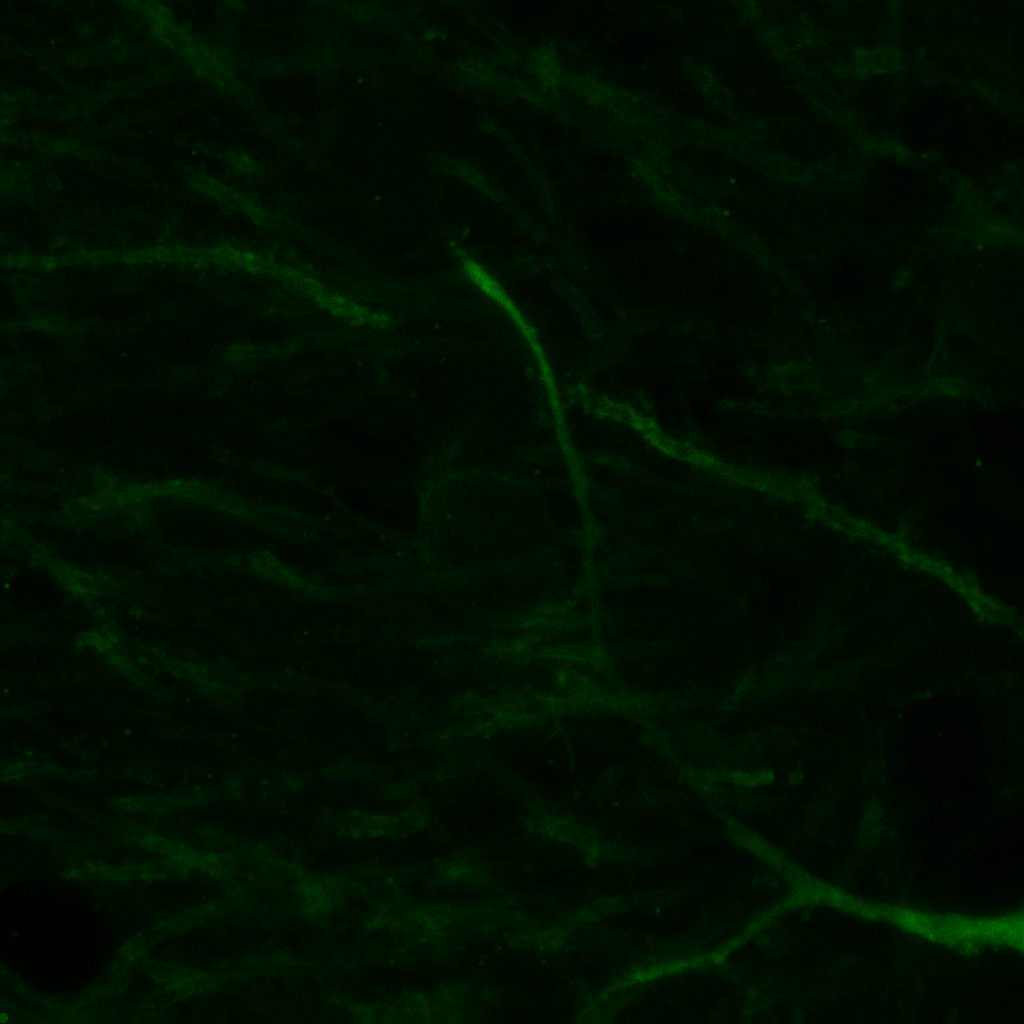

### 3.bmp

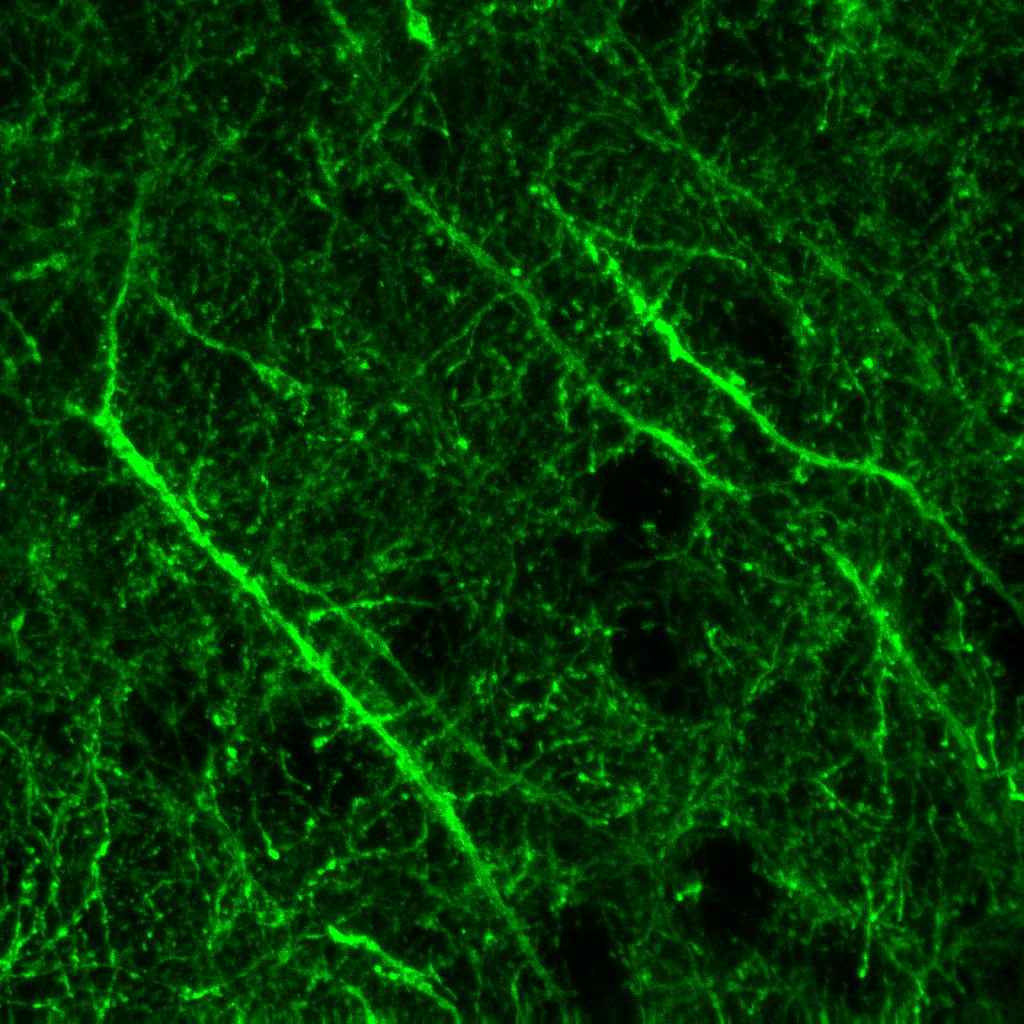

### 3.bmp

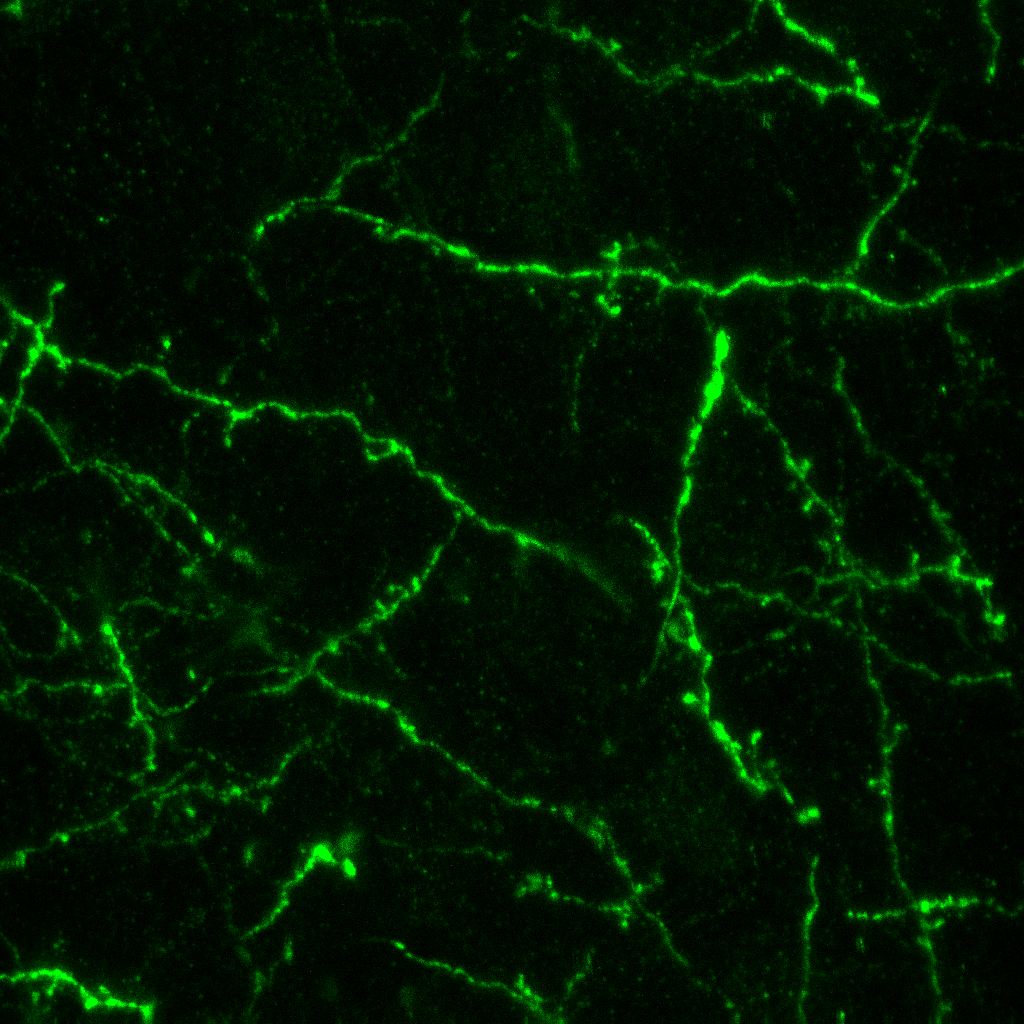

### 3.bmp

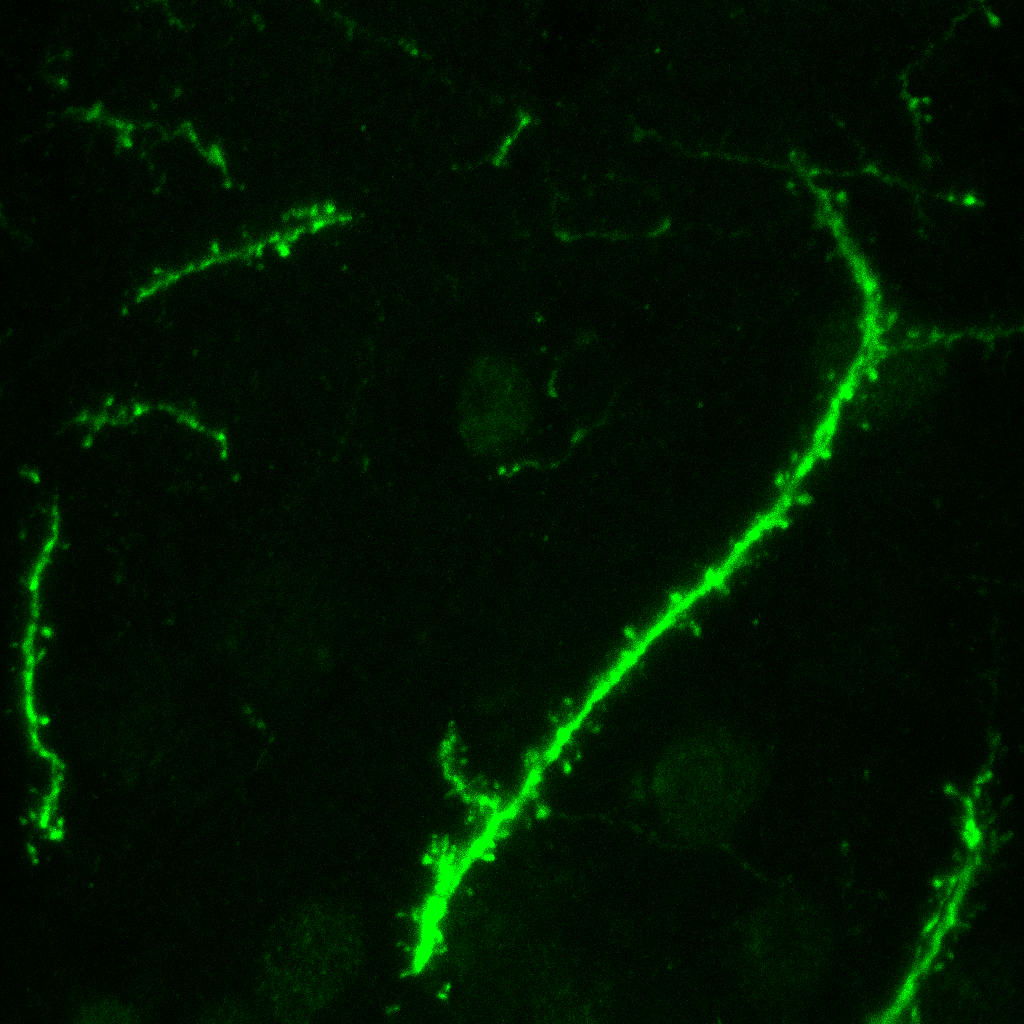

### 3.bmp

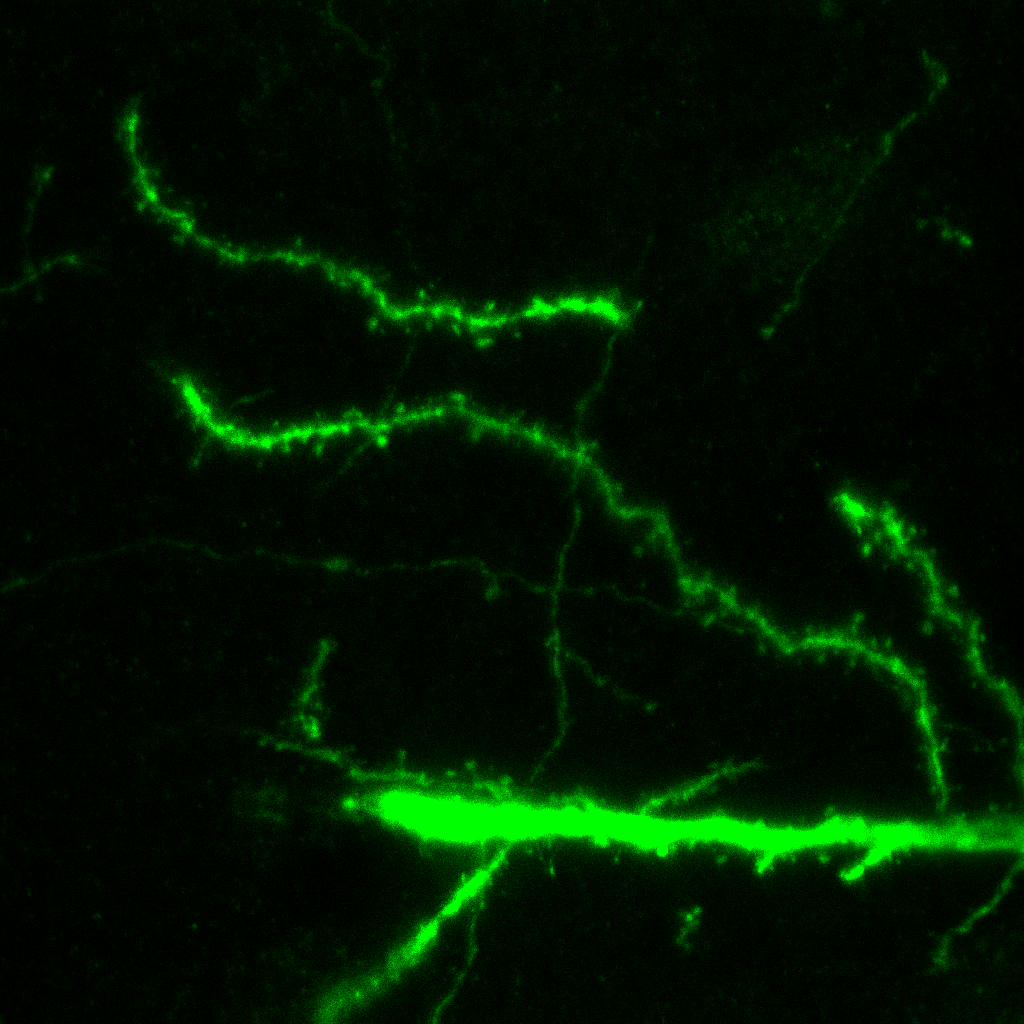

### 3.jpg

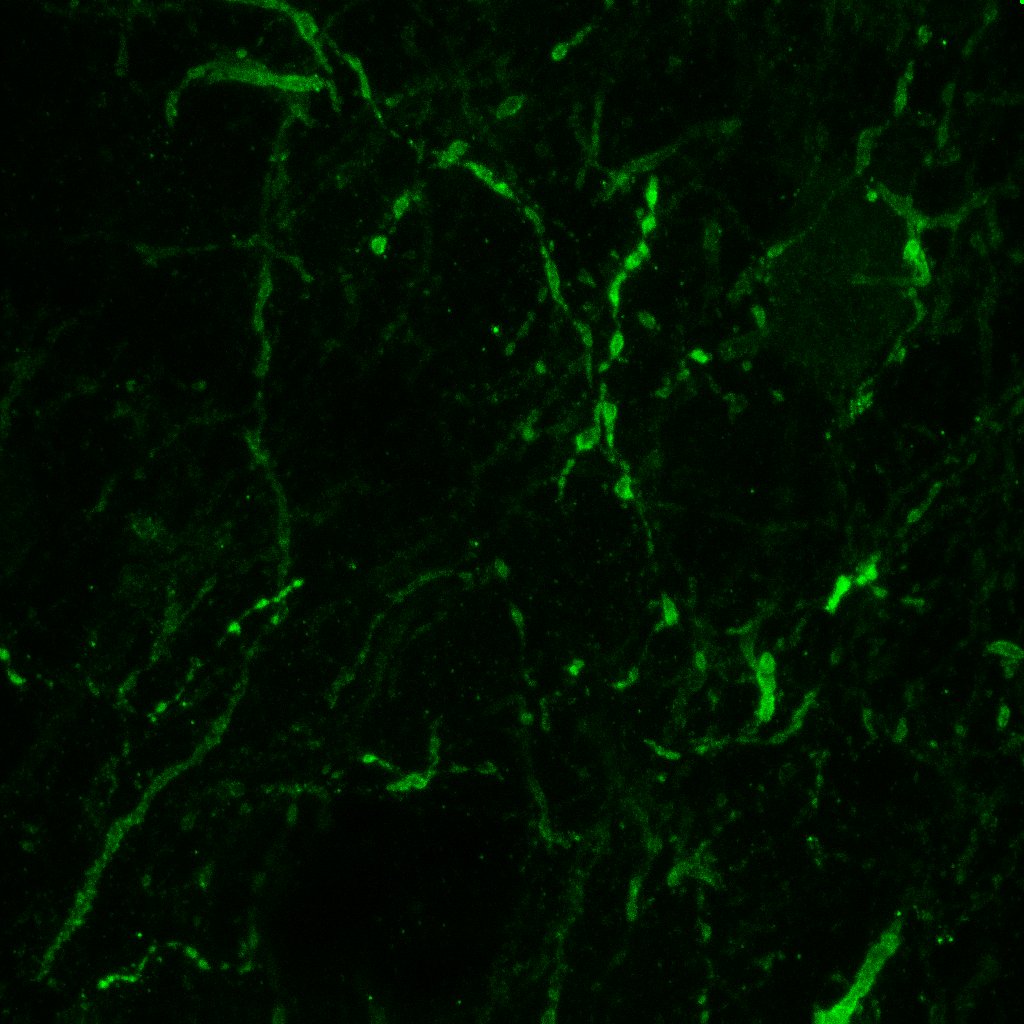

### 3.jpg

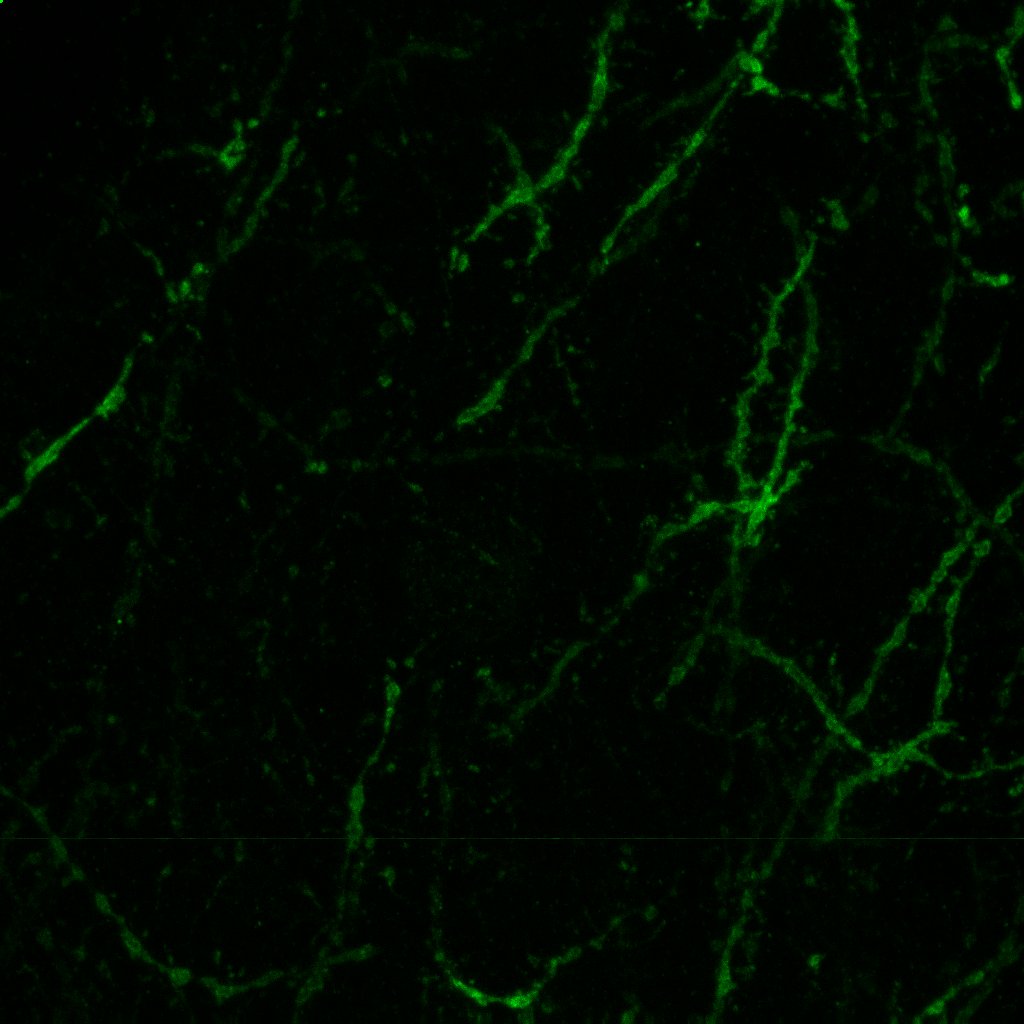

### 3.jpg

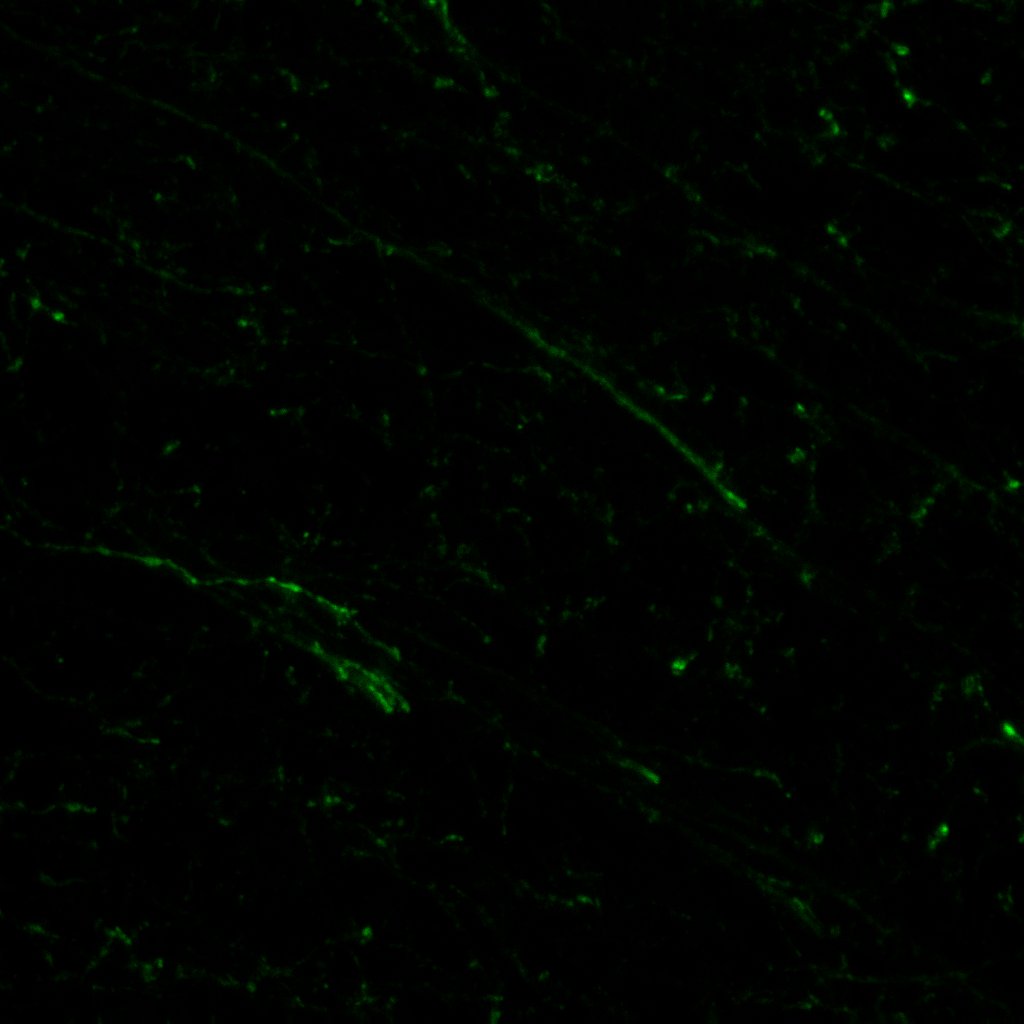

### 3.jpg

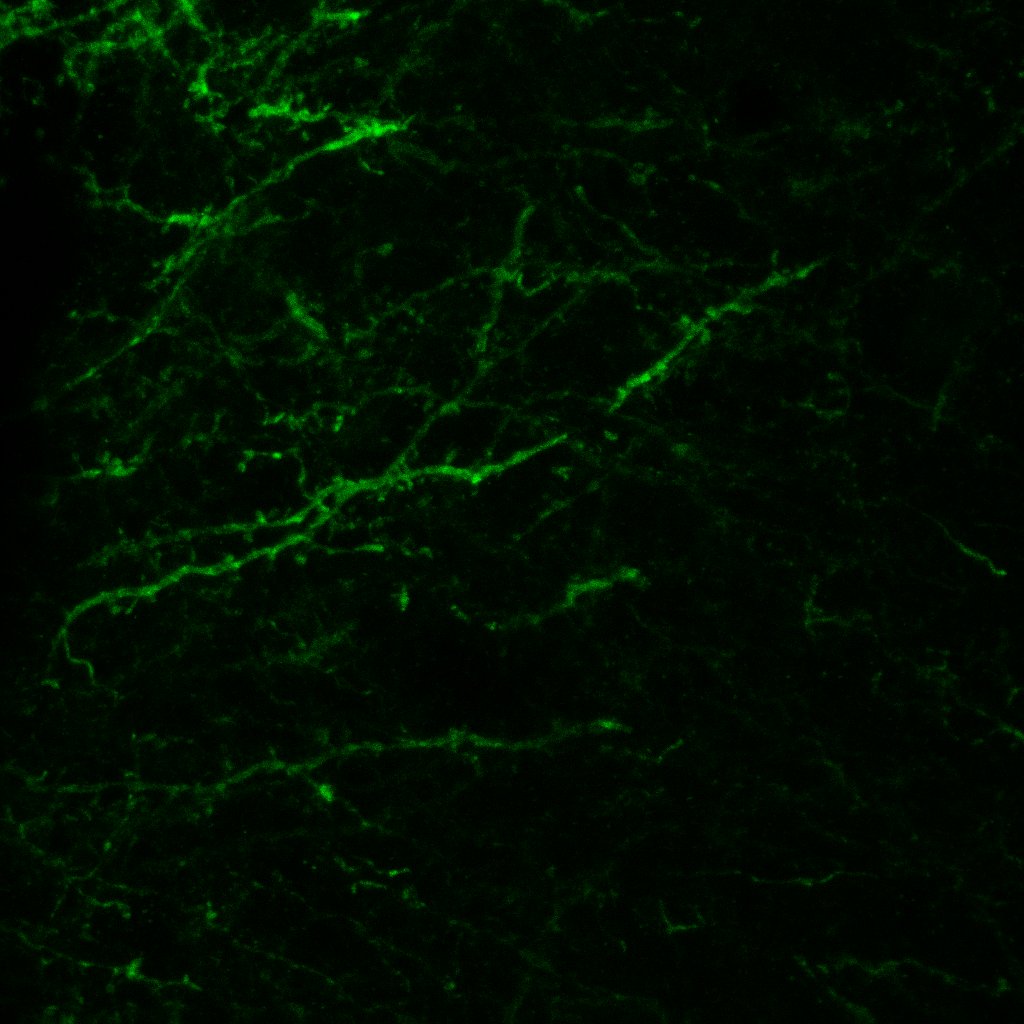

### 3.jpg

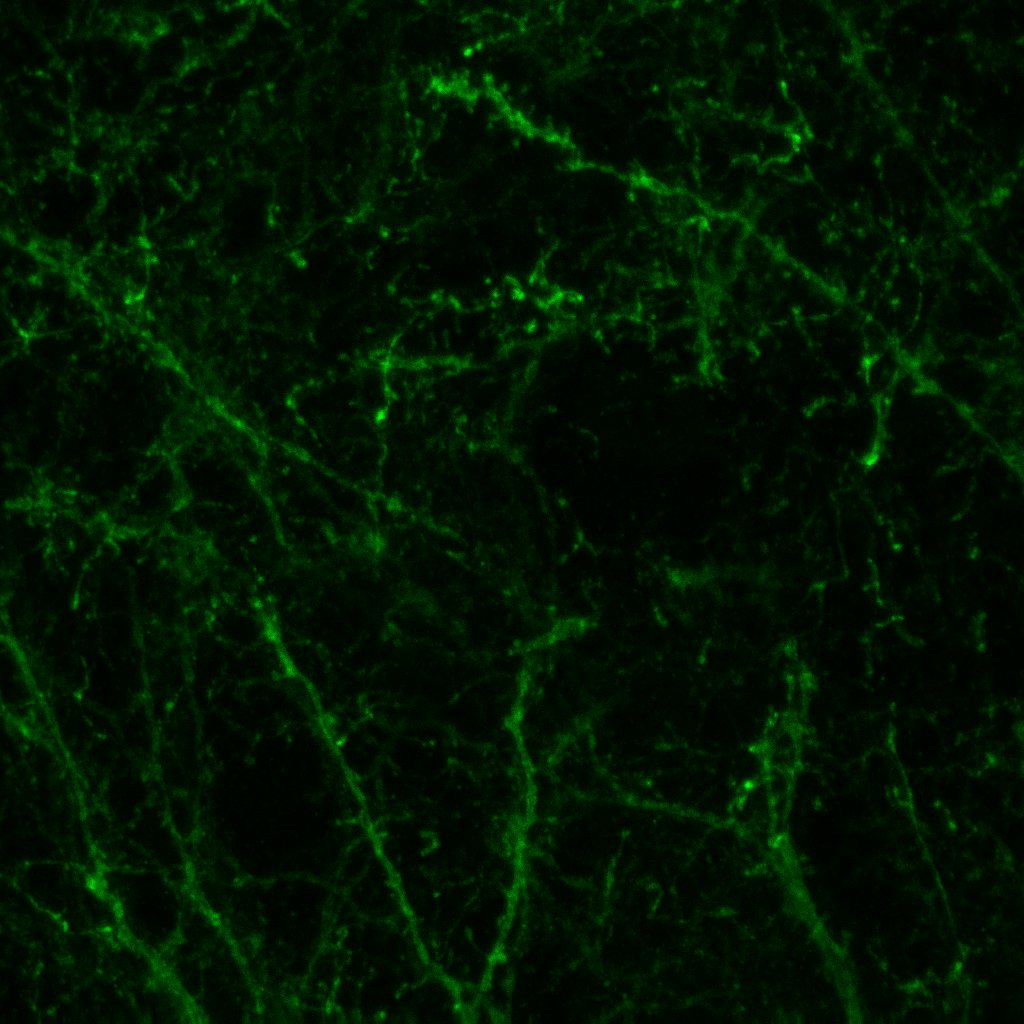

### 3.jpg

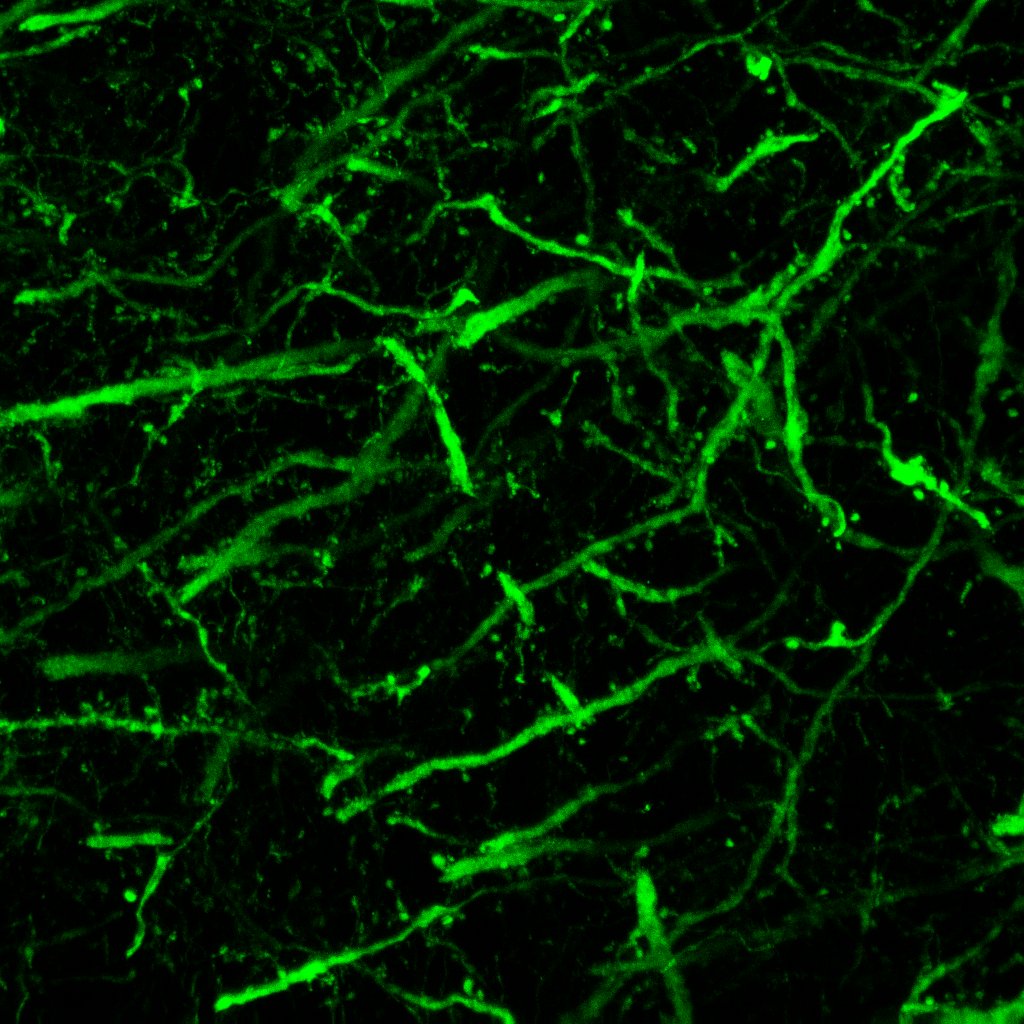

### 3.jpg

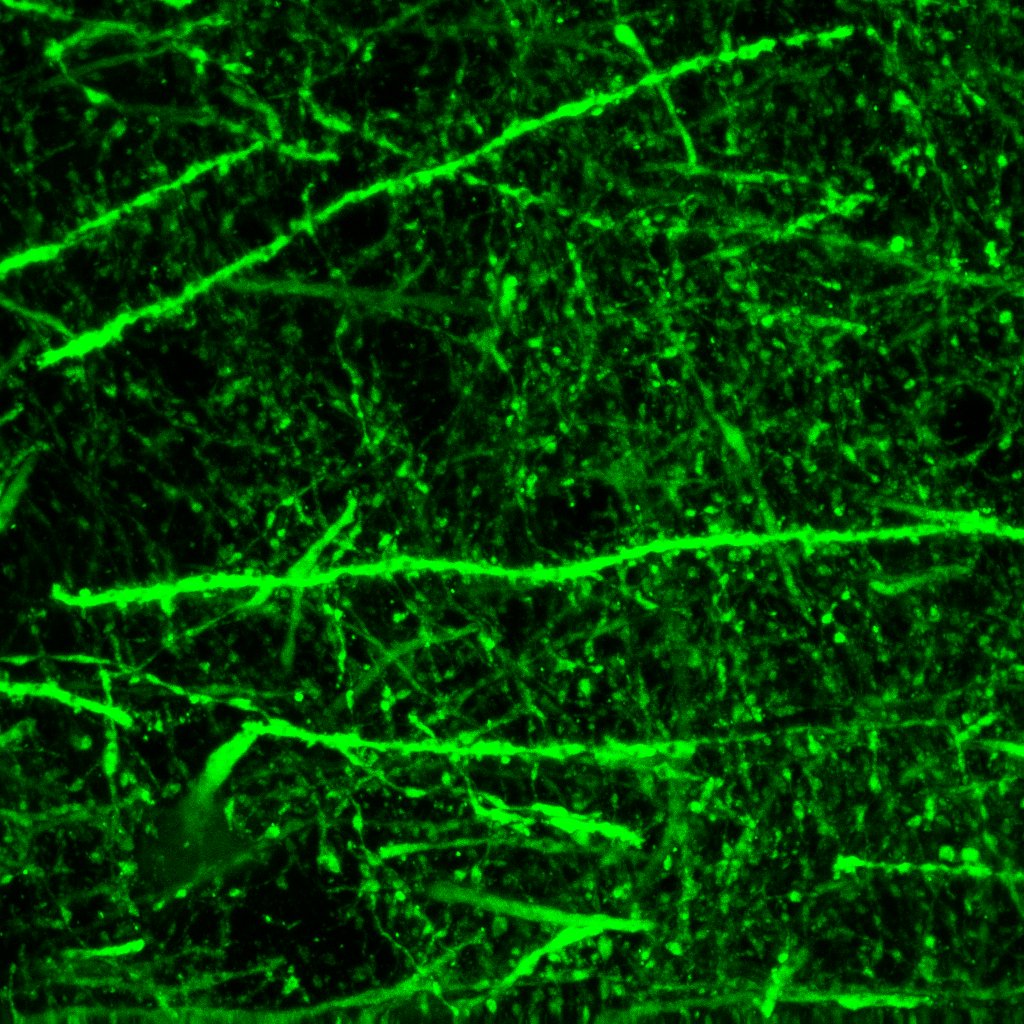

### 4.bmp

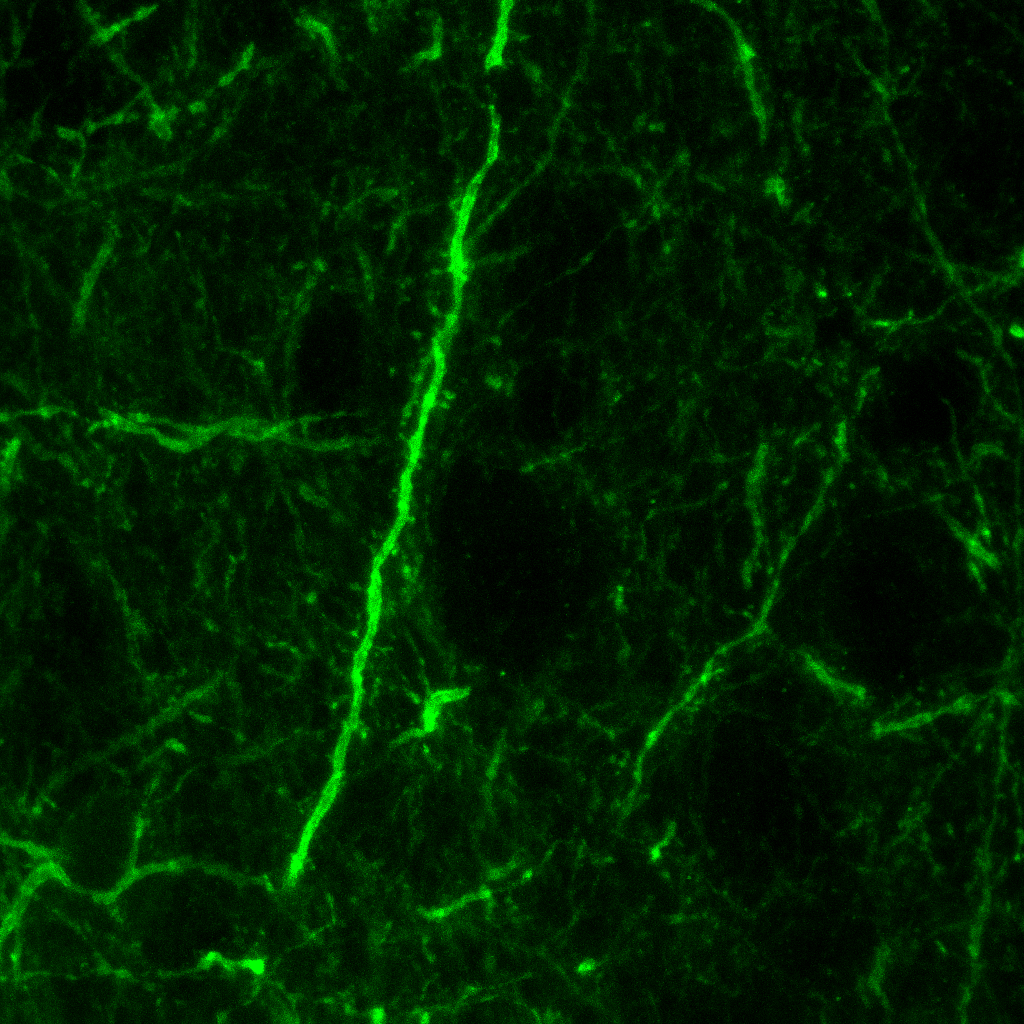

### 4.bmp

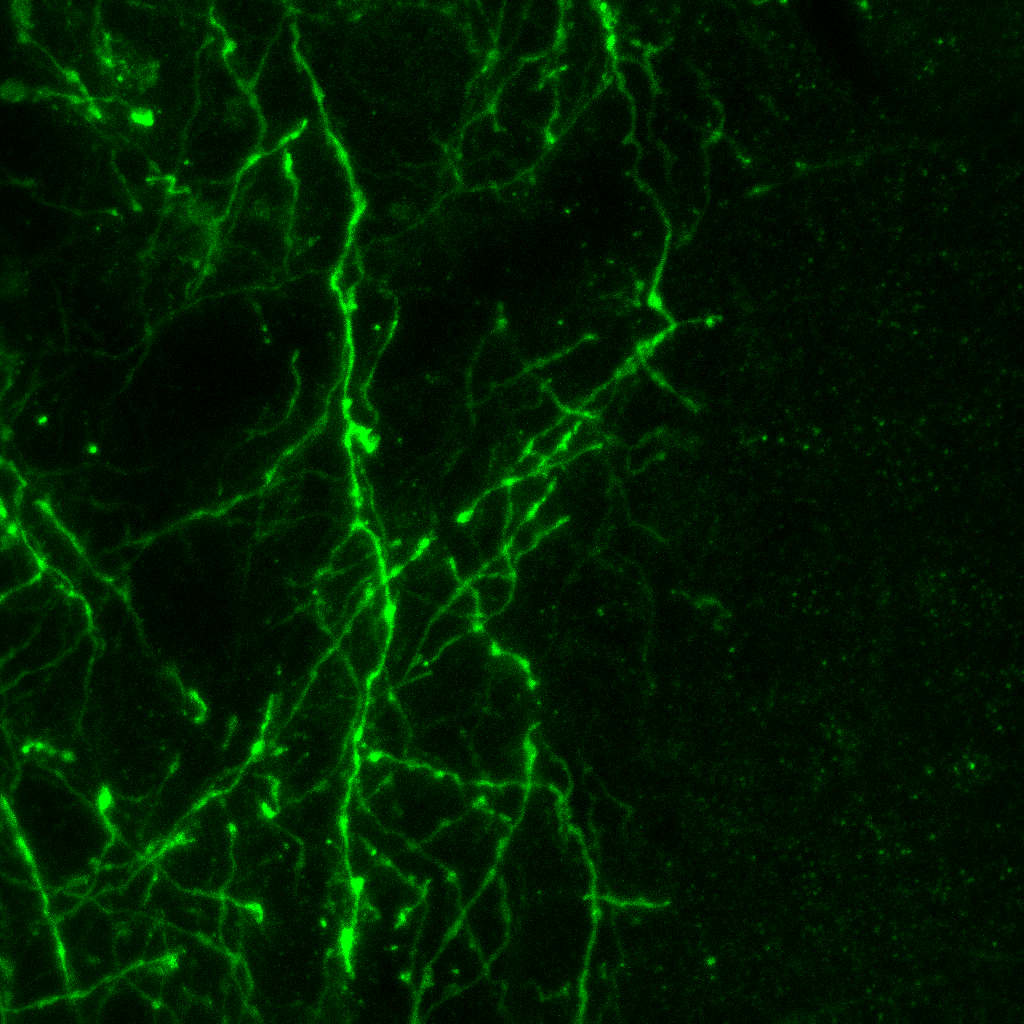

### 4.bmp

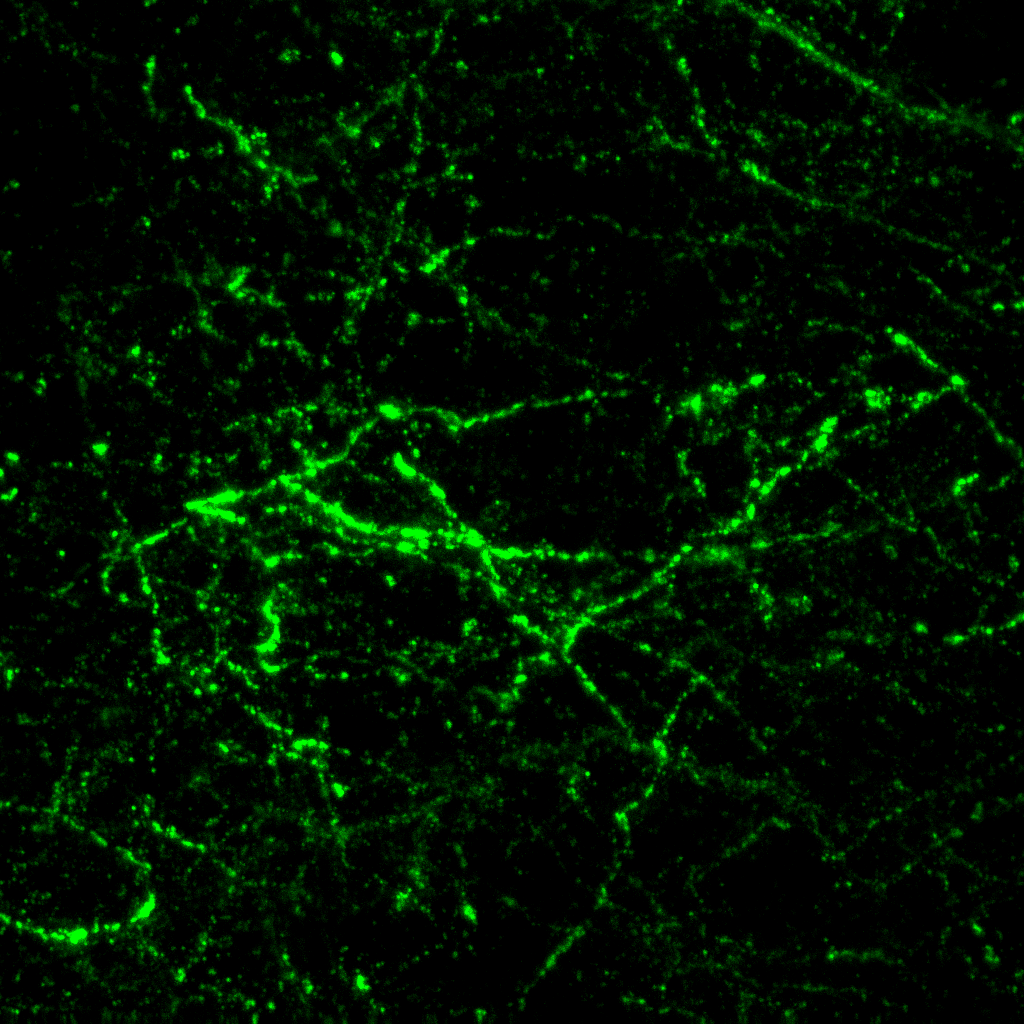

### 4.bmp

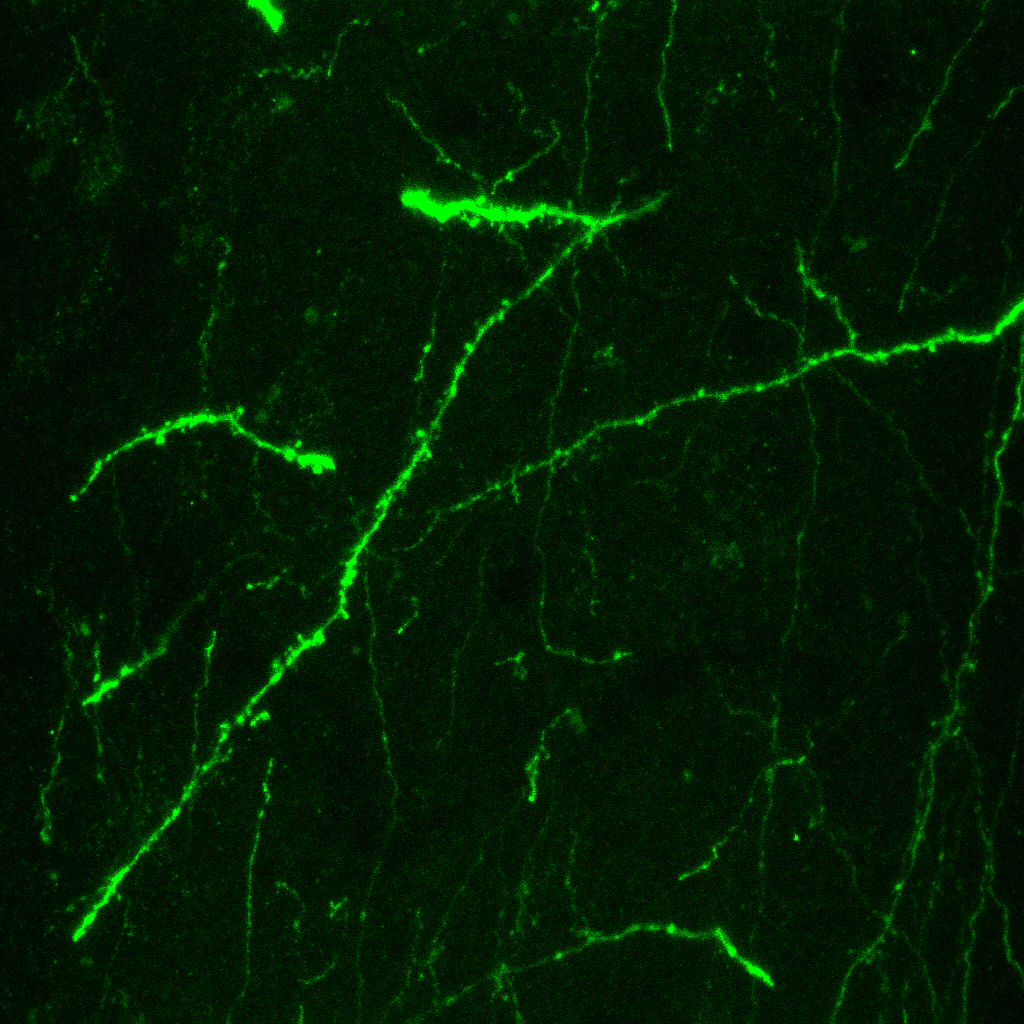

### 4.bmp

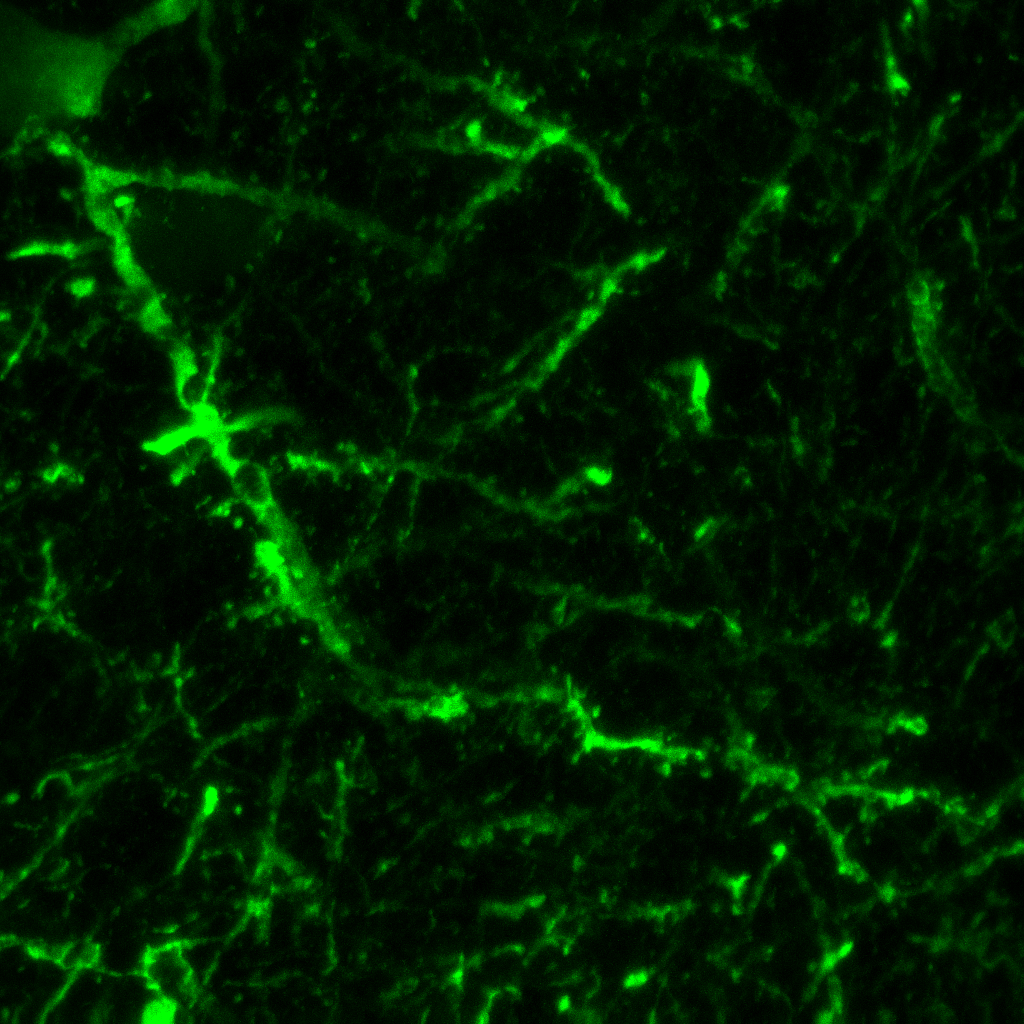

### 4.jpg

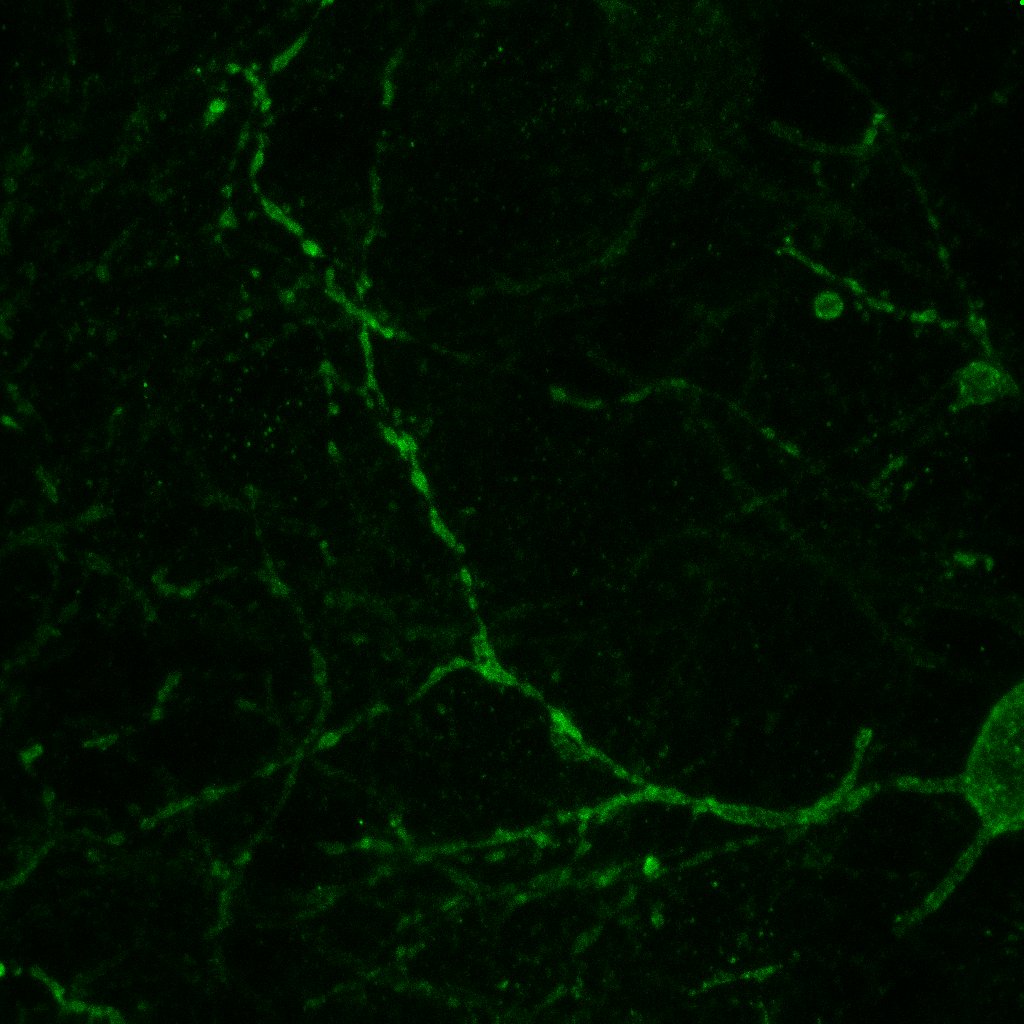

### 4.jpg

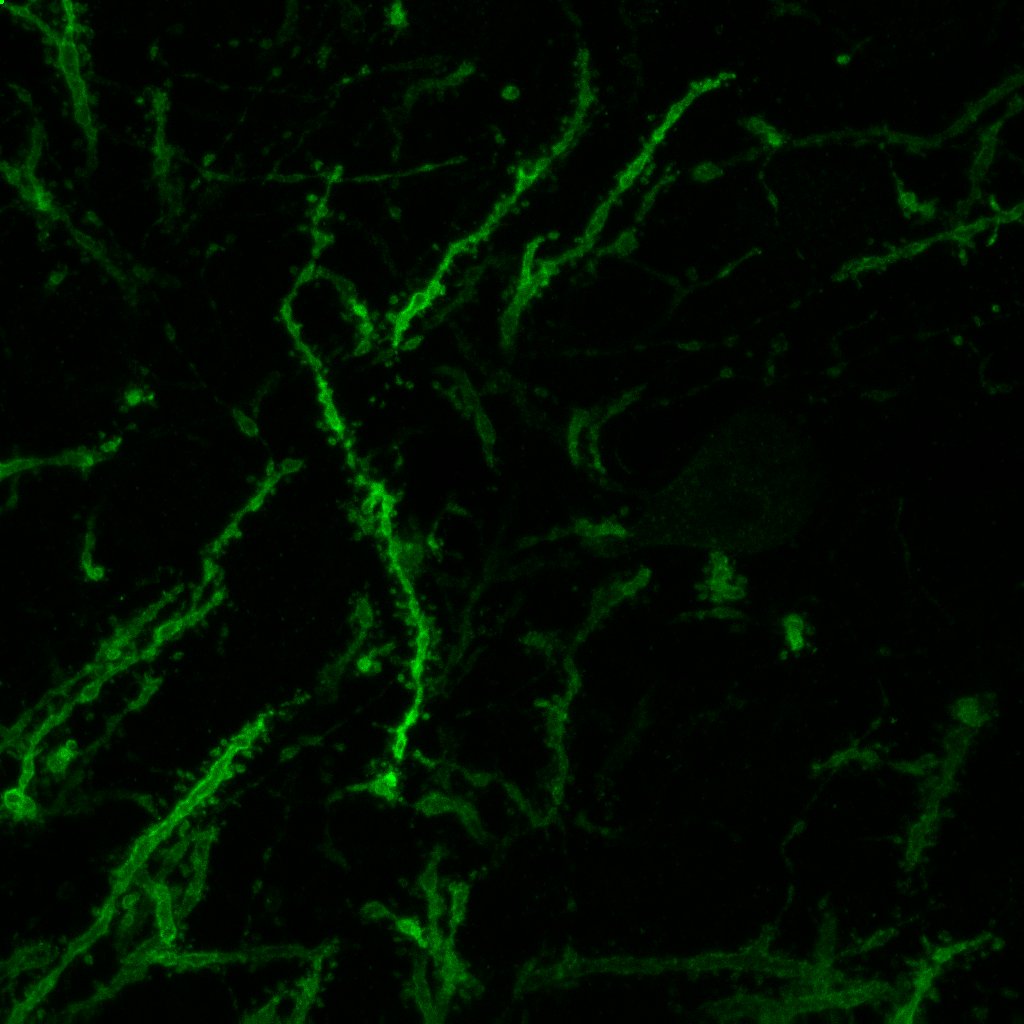

### 4.jpg

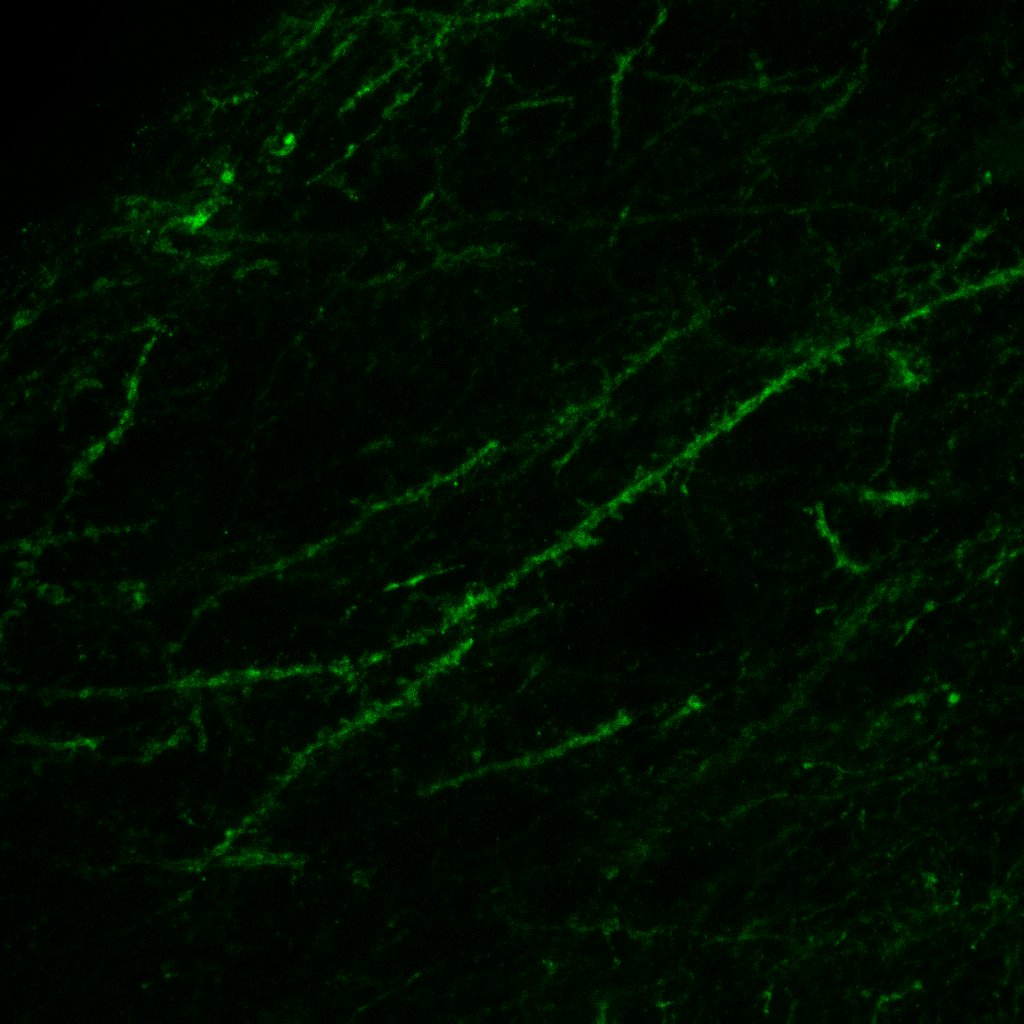

### 4.jpg

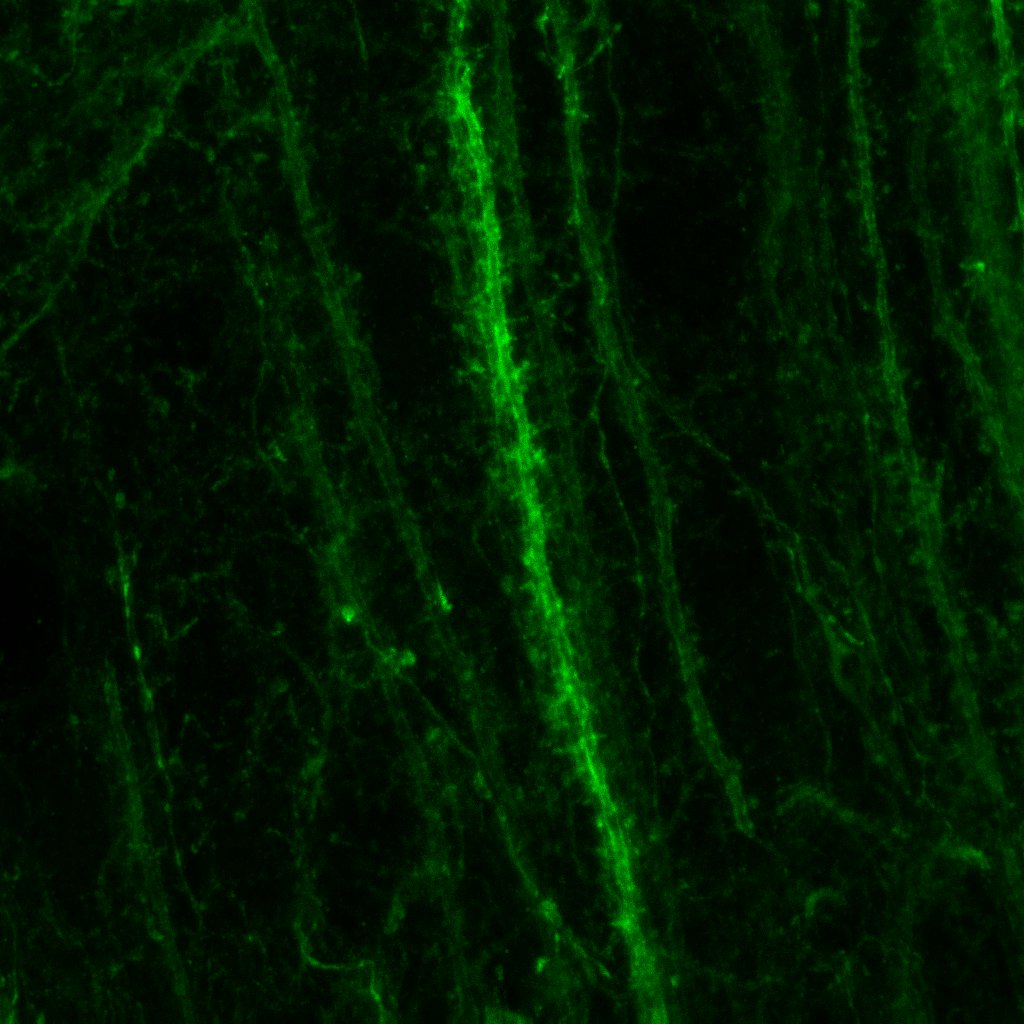

### 4.jpg

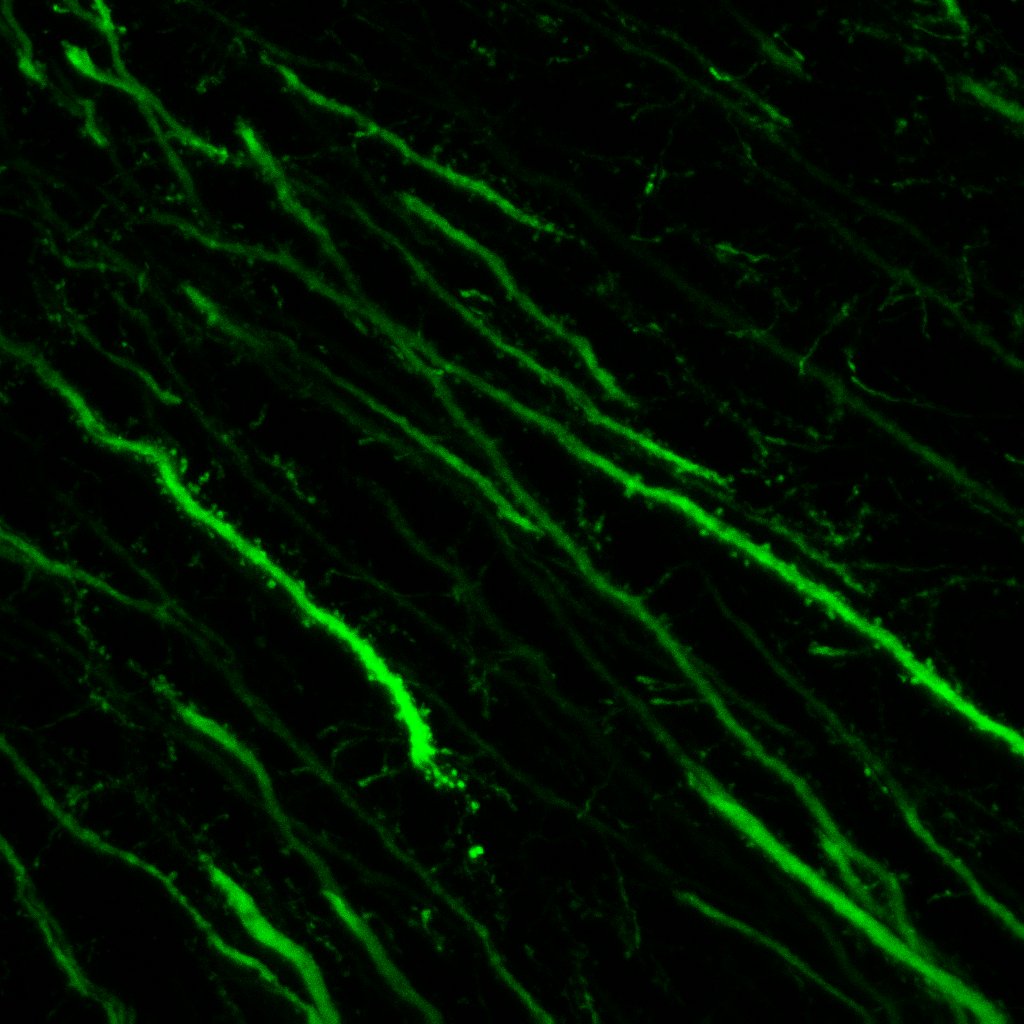

### 4.jpg

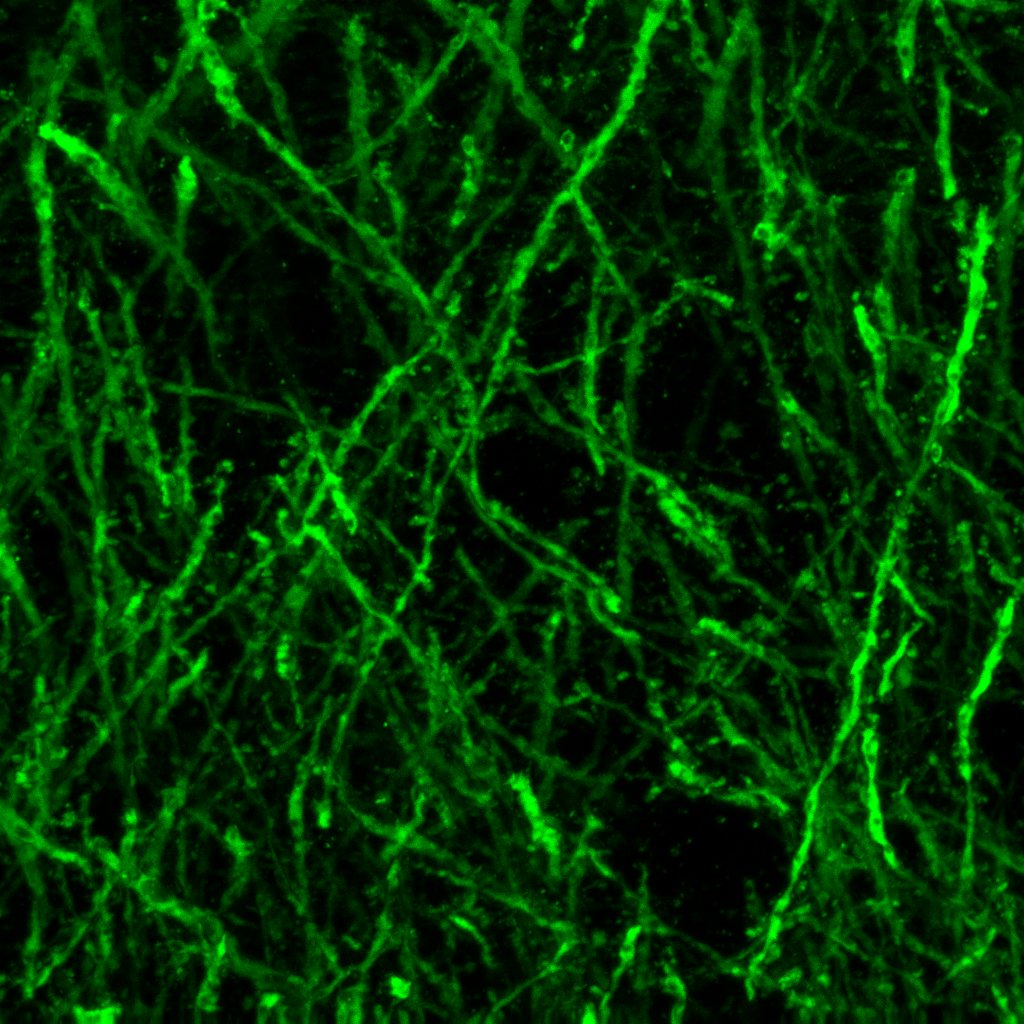
